# Multi-pass, single-molecule nanopore reading of long protein strands with single-amino acid sensitivity

**DOI:** 10.1101/2023.10.19.563182

**Authors:** Keisuke Motone, Daphne Kontogiorgos-Heintz, Jasmine Wee, Kyoko Kurihara, Sangbeom Yang, Gwendolin Roote, Yishu Fang, Nicolas Cardozo, Jeff Nivala

**Affiliations:** Paul. G. Allen School of Computer Science and Engineering, University of Washington, Seattle, WA, USA; Molecular Engineering and Science Institute, University of Washington, Seattle, WA, USA

## Abstract

The ability to sequence single protein molecules in their native, full-length form would enable a more comprehensive understanding of proteomic diversity. Current technologies, however, are limited in achieving this goal. Here, we establish a method for long-range, single-molecule reading of intact protein strands on a commercial nanopore sensor array. By using the ClpX unfoldase to ratchet proteins through a CsgG nanopore, we achieve single-amino acid level sensitivity, enabling sequencing of combinations of amino acid substitutions across long protein strands. For greater sequencing accuracy, we demonstrate the ability to reread individual protein molecules, spanning hundreds of amino acids in length, multiple times, and explore the potential for high accuracy protein barcode sequencing. Further, we develop a biophysical model that can simulate raw nanopore signals *a priori,* based on amino acid volume and charge, enhancing the interpretation of raw signal data. Finally, we apply these methods to examine intact, folded protein domains for complete end-to-end analysis. These results provide proof-of-concept for a platform that has the potential to identify and characterize full-length proteoforms at single-molecule resolution.

---

Annotating the complexity of protein variation is important in understanding biological processes, identifying disease states, and developing effective therapeutics. Proteoform diversity is a term that refers to the vast array of protein variations that can exist due to differences in transcription, translation, and post-translational modifications (PTMs) which can occur through enzymatic (e.g. phosphorylation) and non-enzymatic (e.g. spontaneous deamidation) processes^1^. These variations occur independently and in combination with each other on single protein molecules, creating a “PTM code” that plays unique and specific roles in driving biological processes^2–4^. The ability to sequence single protein molecules in their natural, full-length state would be valuable for better understanding this proteoform diversity and its underlying code. However, current protein sequencing and fingerprinting methods, including Edman degradation and mass spectrometry, have difficulty analyzing full-length proteins from complex samples and face challenges with detection sensitivity, dynamic range, analytical throughput, and instrumentation cost^5,6^. To address these challenges, complementary or potentially disruptive platforms for next-generation protein analysis and sequencing have been proposed, including single-molecule fluorescence labeling and affinity-based approaches^7–10^. However, these emerging techniques also face limitations compared to nanopore technology^11^, which has the potential to achieve direct, label-free, full-length protein sequencing^12^.

Nanopore technology consists of a nanometer-sized pore within an insulating membrane that separates two electrolyte-filled wells^13^. A voltage applied across the membrane drives ionic current flow through the nanopore sensor. When individual analyte molecules pass through the pore, they can manifest a detectable signal change. This change can provide insight into the molecular nature of the analyte. Although originally envisioned and now commercialized as a technique for sequencing nucleic acid strands, nanopore sensing has great potential for protein analysis^13,14^. It has been used for discrimination of peptides and proteins^15–22^, real-time measurement of protein-protein^23^ and protein-ligand interactions^24^, and aptamer-mediated protein detection^24,25^. Additionally, protein nanopores have shown promise in identifying amino acids and PTMs, such as phosphorylation and glycosylation^26,27^, which serve as important biomarkers of cell states and diseases. Recent work has demonstrated some ability to read DNA-conjugated peptide strands using DNA-processive molecular motors, such as a helicase or polymerase^28–30^. Further, rereading of peptide fragments using this strategy made it possible to resolve amongst a small subset of single-amino acid substitutions with high accuracy^29^. Despite this progress, obtaining sequence information from intact, full-length protein strands using nanopores has been hindered by the difficulty in driving long protein strands through the sensor, which arises due to the neutrally charged polypeptide backbone, varying charge states of amino acid side chains, and stable tertiary structures^31^.

To overcome the challenges of reading full-length protein molecules, here, we developed a technique to reversibly thread long protein strands into a CsgG pore^32^ by electrophoresis, and then enzymatically pull them back out of the pore using the protein unfoldase/translocase activity of ClpX^33^. While the initial stage of threading the protein into the pore using electrophoretic force happens too quickly to resolve any sequence attributes, unfoldase-mediated translocation of proteins back out of the pore manifests slow, reproducible ionic current signals. This method enabled the processive translocation of long proteins, facilitating the detection of numerous single amino acid substitutions and PTMs across protein strands up to hundreds of amino acids in length. We also developed an approach to rereading the same protein strand multiple times. Furthermore, we show this method enables unfolding and translocation of entire folded protein domains for linear, end-to-end analysis.

## An unfoldase-mediated approach to nanopore protein sequencing on a nanopore array

While we and others have previously developed approaches to unfoldase-mediated protein translocation through nanopores^34–36^, these methods required complex experimental setups and did not demonstrate the level of single-amino acid sensitivity required for sequencing. In our previous methodology, the unfoldase and its cofactors were located in the *trans*-side solution, opposite to the location of the protein substrate addition^34,35^. This setup rendered the method incompatible with commercial high-throughput nanopore sensor array devices, such as Oxford Nanopore Technologies’ MinION, which do not allow access to the *trans* compartment solution. To overcome the need for *trans* motor addition, we designed a more streamlined two-step process. First, the protein substrate is threaded into the nanopore via electrophoretic force (*cis*-to-*trans*). Then, ClpX is added to the *cis* solution to steadily pull the substrate protein back out of the pore (*trans*-to-*cis*) (**Fig. 1a**).

**Fig. 1.**
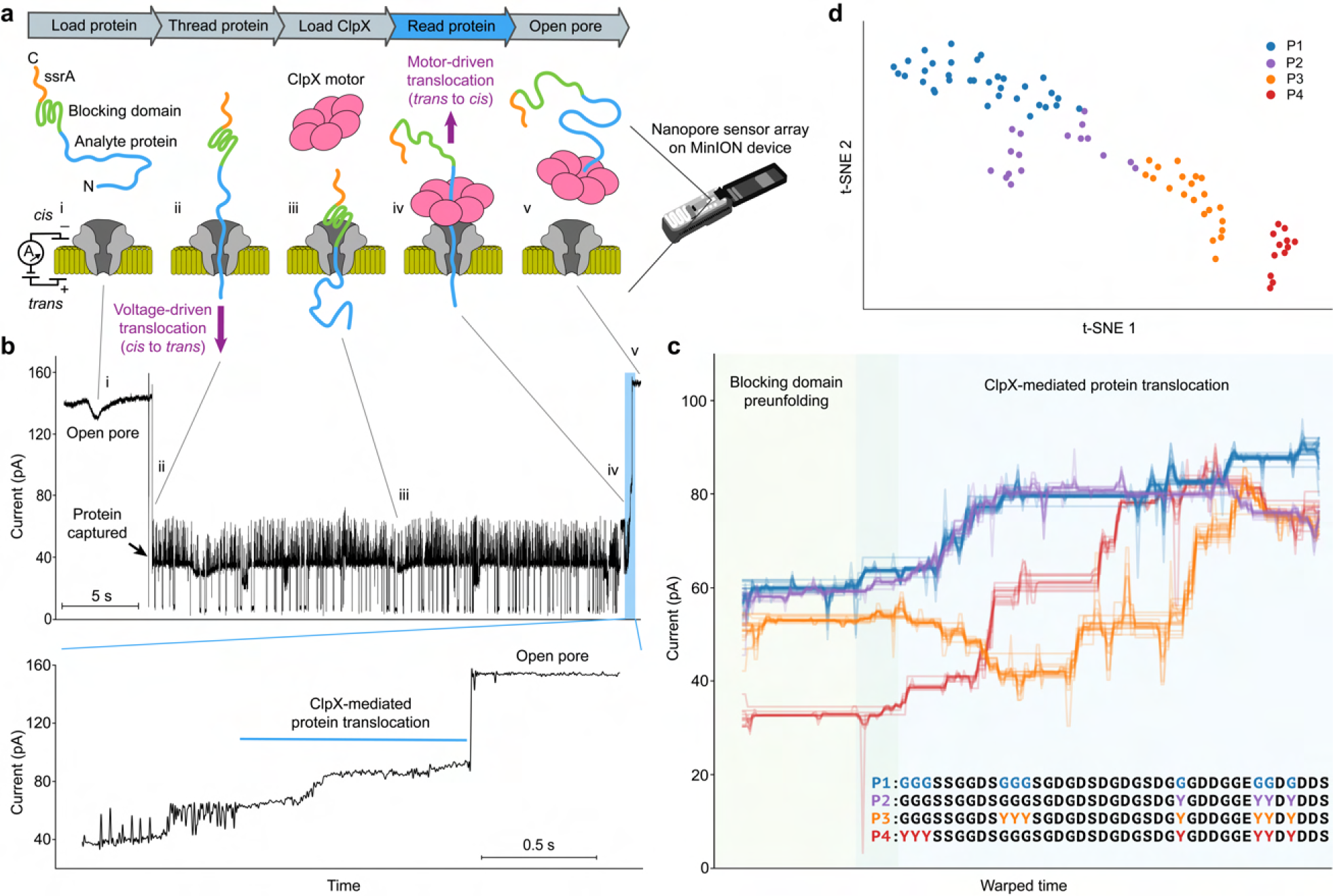
Nanopore protein reading using an unfoldase. **a**, Schematic of *cis*-based unfoldase approach on the MinION platform. Assigned Roman numerals correspond to ionic current states in **b. b**, Example trace of protein P1. **c**, Ensemble traces of protein P1 (blue, *n* = 34) and mutants P2 (purple, *n* = 17), P3 (orange, *n* = 21), and P4 (red, *n* = 12). Protein sequences are oriented from C to N, with all mutation regions shown in color. **d**, t-Distributed stochastic neighbor-embedding (t-SNE) plot derived from embedding the all pairwise signal DTW distance comparison matrix (**Methods**).

We first synthesized a protein to evaluate this method, which comprised an unstructured N-terminal sequence of 42 amino acids rich in glycine, serine, and aspartic acid (polyGSD), attached to a stably folded domain (Smt3). This was followed by a short, positively charged sequence (RGS repeat) and a ClpX-binding ssrA tag at the C-terminal end (referred to as protein P1; **Supplementary Fig. 1**). The combination of the RGS and the folded domain was included in the design to inhibit complete translocation of the protein through the pore, thereby preserving the ssrA tag’s accessibility in the cis compartment. After introducing P1 into a MinION R9.4.1 flow cell, which incorporates a CsgG pore variant (Oxford Nanopore Technologies)^37^, and applying a voltage of −180 mV, we observed current blockades associated with the capture of the negatively-charged protein tail within the pores. To ascertain whether ClpX could be used to extract the captured protein from the nanopore, we then introduced a buffer solution containing ClpX and ATP into the flow cell. Under these conditions, we observed events in which the deep ionic current blockade, characteristic of capture of the substrate protein in the nanopore, returned to the open channel state in a near stepwise manner sometime after ClpX addition (**Fig. 1b** and **Supplementary Fig. 2**). We also determined that these events were ATP-dependent and occurred at a slower rate in the presence of ATPγS (**Supplementary Fig. 3**), an ATP analog that is more difficult for ClpX to hydrolyze^38^. These results are consistent with our model that ClpX was binding to the ssrA tag and translocating the captured protein out of the nanopore in the C to N terminal directionality.

If this was true, we reasoned that mutations in the tail domain of the protein would induce alterations in the ionic current states observed during ClpX-mediated translocation of the protein through the nanopore. In order to test this, we synthesized three new proteins (P2, P3, and P4), each containing several tyrosine (Y) mutations at distinct positions of the polyGSD sequence (**Supplementary Fig. 1**). **Fig. 1c** shows ensemble ionic current traces for each of these proteins (individual traces in **Supplementary Fig. 2**). To directly compare the signal profiles of the four protein sequences, we used the Dynamic Time Warping (DTW) Barycenter Averaging algorithm^39^, which aligns and aggregates the raw signals. This revealed that the major differences across the translocation signals corresponded with the respective positions of the tyrosine mutations along the different protein strands (**Fig. 1c**). Moreover, analysis by all-vs-all signal comparison using DTW and t-SNE dimensionality reduction (**Methods**) showed that the sets of translocation signals generated by each unique protein sequence were dissimilar (**Fig. 1d**). These results demonstrated that amino sequence-dependent signal features could be observed using this approach.

## Resolving single-amino acid substitutions

Having established the *cis*-based ClpX approach, we next sought to investigate the sensitivity of this method to single-amino acids as a first step towards developing a long-read protein analysis method. To do this, we designed protein constructs that included five repeating sequence blocks, each containing 59 amino acids. These blocks were built with a base sequence of glycine, serine, aspartic acid, and glutamic acid (**Fig. 2a** and **Supplementary Fig. 1**). We introduced a unique amino acid mutation at the central position within each block and demarcated the blocks with a double tyrosine mutation at each end. This design was used to minimize potential overlapping signal contributions from the mutations by ensuring sufficient spacing to prevent concurrent pore occupancy by multiple mutations. This hypothesis was grounded on prior observations indicating that around 20 amino acids can occupy the CsgG sensing region when in a stretched conformation^40^. We termed these strategically designed protein constructs as Proteins for Amino acid Sequencing Through Optimized Regions (PASTORs). We synthesized a total of eight different PASTOR variants, each containing a different sequence of mutations. The PASTOR design allowed us to analyze up to five different mutations in a single nanopore read, and the total set of eight pastors allowed us to interrogate each of the 20 amino acids in two different PASTOR sequence contexts (C-to-N): HDKER, GNQST, FYWCP, AVLIM, VGDNY, TWAFH, PRMQE, and KSILC.

**Fig. 2.**
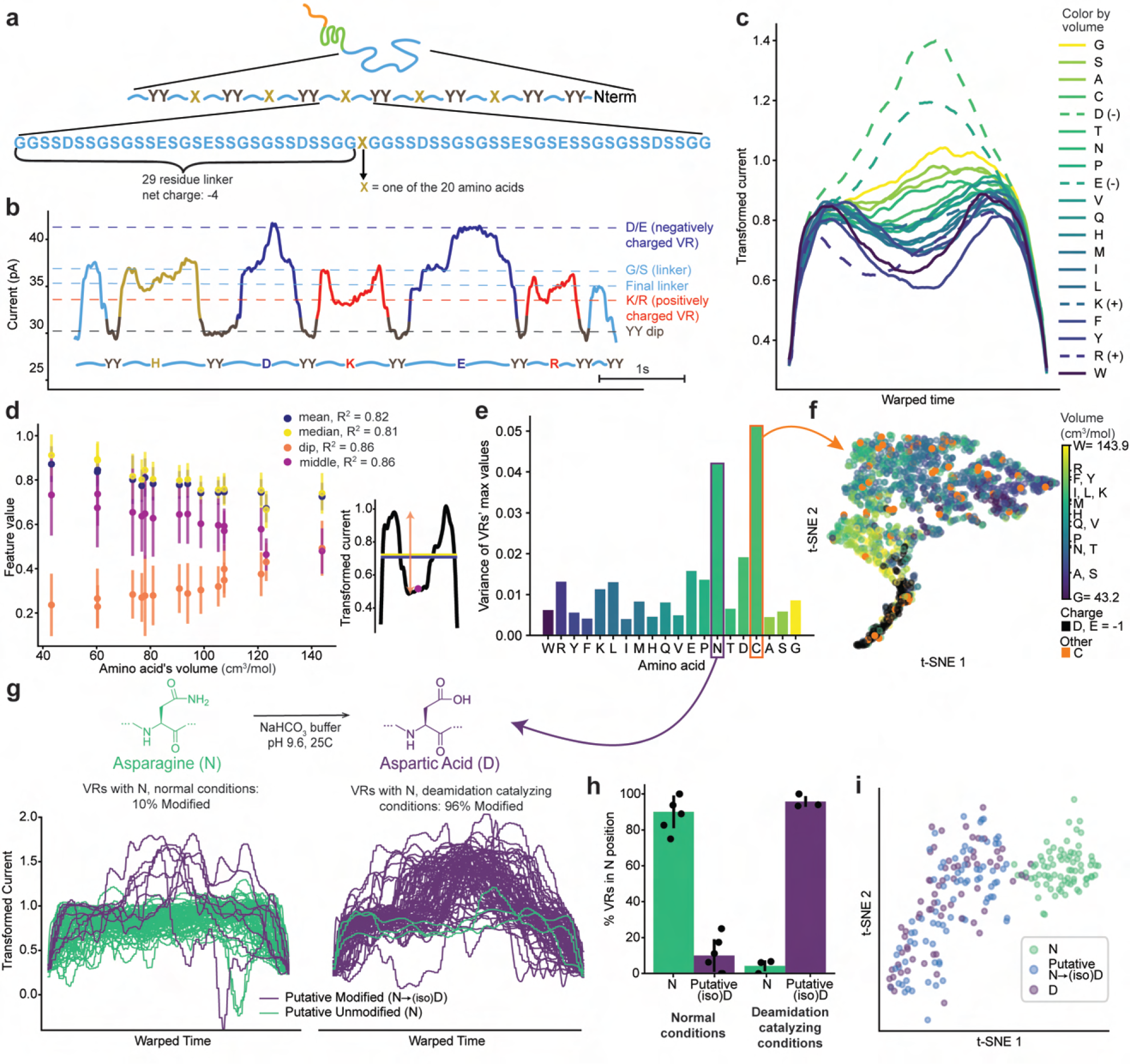
Detecting single amino acid mutations across long protein strands. **a**, PASTOR sequence composition. **b**, Filtered nanopore current trace of PASTOR-HDKER. Regions’ color boundaries are defined by YY-segmentation. **c**, Average signal trace for each amino acid’s transformed VRs, after euclidian alignment of all the VRs equidistantly stretched to the same length. VRs corresponding to a charged amino acid are shown in a dashed line. **d**, Scatter plot of various features of the VRs, with error bars denoting one standard deviation and explanation of the features to the right. *n* varies from 56 to 98. **e**, Bar blot of the variance of the max value of the transformed VRs corresponding to each amino acid. **f**, t-SNE map showing clustering of the pairwise DTW distance between each amino acid, with all amino acids other than D, E, and C being colored by the volume, the negative amino acids colored black, and C being highlighted in orange. **g**, Plot of all the VRs corresponding to asparagine in normal conditions (left) and in conditions that catalyze the deamidation of asparagine to aspartate (right). Lines colored teal if the max value of the transformed signal < 1.3, and purple otherwise. *n* = 81 for normal conditions and 77 for deamidation conditions. **h**, Bar plot displaying percent of mutations that have been putatively deamidated or not (same threshold as in g, h) in VRs corresponding to asparagine, across technical replicates with *n* = 6, 4, 3, and 3 from left to right. Error bars denote standard deviations. **i**, t-SNE plot as in g, showing only asparagine and aspartate VRs. Asparagine VRs are colored purple if the max value of the transformed signal < 1.3, and blue otherwise.

ClpX-mediated analysis of the PASTOR proteins manifested elongated ionic current traces containing repetitive patterns that resulted from the seven YY mutations, seen as seven repeated dips in the signal preceding return to open channel (PASTOR-HDKER is shown in **Fig. 2b** and additional PASTORs in **Supplementary Fig. 4**). Between these dips, distinctive and reproducible variations in the ionic current signals were observed, corresponding with the variable amino acid mutation within each block. Utilizing the consistent, substantial effect of the YY mutations, we segregated the current signals into regions termed ‘YY dips’ and ‘variable regions’ (VRs) (**Supplementary Figs. 5 and 6**). This approach allowed us to analyze the unique ionic signatures of each amino acid mutation in isolation. We found that the ionic current levels of variable regions (VRs) with a neutral amino acid mutation showed a negative correlation with the volume of the amino acid (**Figs. 2c** and **2d**). This observation reinforces a volume exclusion model, where larger amino acids block more current than their smaller counterparts. Interestingly, the VRs containing positively charged residues (K/R) decreased the current level below the baseline sequence, while negatively charged residues increased it, diverging from the volume exclusion model. This effect was more substantial for negatively charged residues than for positively charged ones. One possible explanation for this could be that the negatively charged residues resist translocation to the negatively charged *cis* compartment, causing the protein strand to stretch and consequently decrease the total volume of protein in the pore. Conversely, a positively charged residue would be attracted to the *cis* compartment and could introduce upstream kinks in the protein strand, adding additional protein volume in the pore. The impact on signal levels could also be attributed to variations in solvation states and the mobility of ions near the charged amino acids. Collectively, these results showed that this method was sensitive to single amino acid residues.

## Detection of a non-enzymatic post-translational modification

Our analyses also revealed substantial read-to-read signal variability within asparagine (N) and cysteine (C) VRs (**Fig. 2e**). The pronounced signal variability associated with cysteine (**Fig. 2f**) is likely due to its capacity to assume a spectrum of oxidation states^41^. Signal traces for asparagine VRs, however, commonly demonstrated one of two distinct signal levels. The majority of signal events presented a minor dip, consistent with the volume exclusion model (**Figs. 2c** and **2g**). Conversely, around 10% of reads, under standard experimental conditions, exhibited a high signal peak, mirroring those of the acidic amino acid VRs aspartate and glutamate. We hypothesized that these deviant reads might arise from post-translational asparagine deamidation, a process which transforms the asparagine sidechain to aspartate or isoaspartate. This conversion is recognized as a common PTM and is speculated to operate as an important biological timing mechanism in-vivo^42^. To investigate this reasoning, we exposed the PASTOR-VGDNY protein to buffer conditions that promote the deamidation of asparagine to (iso)aspartate^43^ (**Methods**). Following protein incubation in this buffer, nanopore analysis of asparagine VRs revealed that approximately 96% now exhibited signal traits similar to aspartate (**Figs. 2g–i** and **Supplementary Figs. 7 and 8**). This increase in aspartate-like signals post-incubation supports our hypothesis that the atypical reads in the asparagine VRs originate from post-translational deamidation.

## Modeling nanopore signals directly from amino acid sequence

Considering the relationship between the volume and charge of individual amino acids and their impact on nanopore signals, we developed a biophysical model designed to simulate nanopore signals from a protein’s amino acid sequence directly. This model, building on Cardozo et al.’s findings^40^, determines a summation of the volume and charge of amino acids within a moving 20-residue window, applying a centrally positioned negative parabolic weight (**Figs. 3a–d**). **Fig. 3e** shows the signal generated by our model for the PASTOR-TWAFH protein sequence aligned to an actual nanopore trace of the same protein. Model signals for all proteins in this study are shown in **Supplementary Fig. 9**. To quantitatively evaluate the congruency between our model and actual ionic current traces, we calculated their average distance post-DTW for a given sequence, denoted as *S*, post-normalizing both the model and experimental current traces. We designated this value as *DTWDS* and compared it to the distribution of DTW distances between the actual ionic current traces and the model of randomly composed sequences of amino acids derived from the distribution of amino acids in *S*, referred to as *R*. The average *DTWDS* for PASTORs ranked in the top 0.2% of the best matches within *R* (**Fig. 3f**). This evaluation affirms that the signal agreement observed in **Fig. 3e** is not due to artifacts from DTW alignment, thereby reinforcing the assertion that our model has the capacity to simulate these current traces accurately within these sequence contexts.

**Fig. 3.**
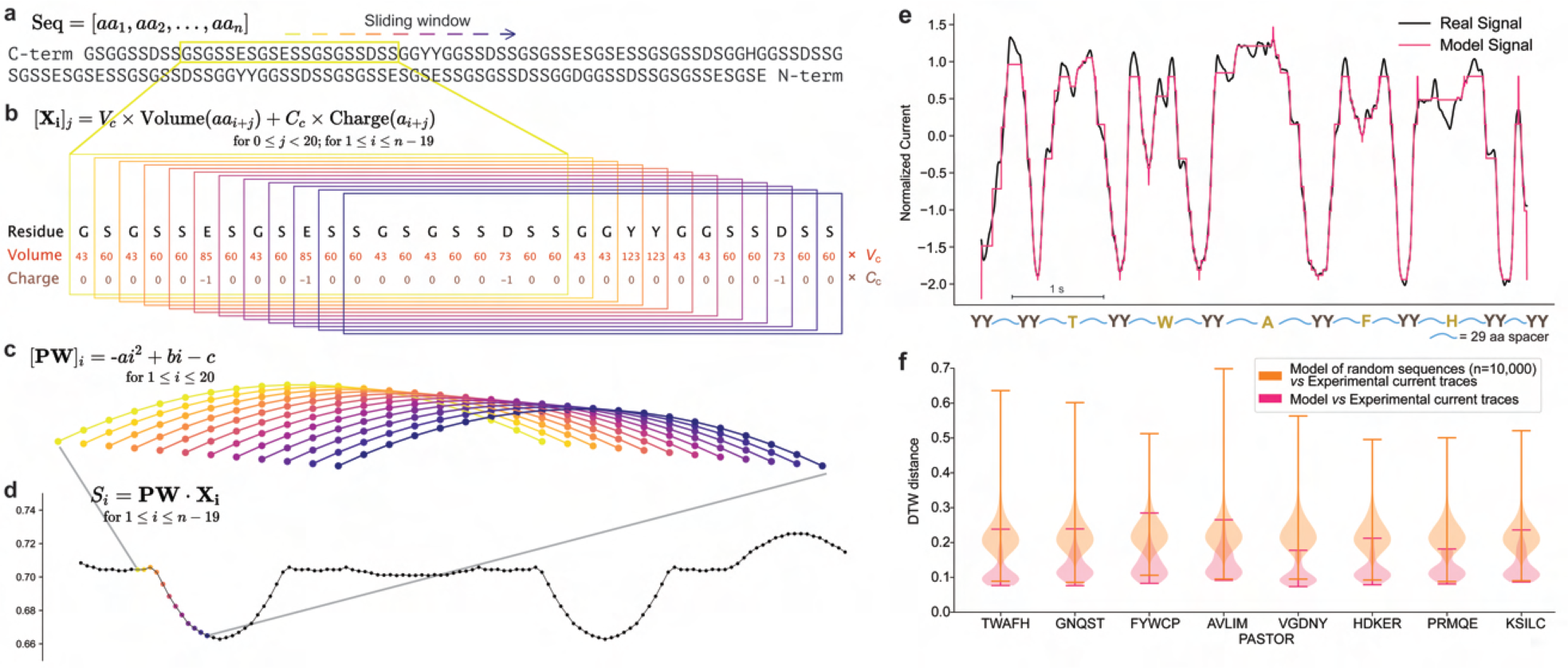
A biophysical model for simulating nanopore ionic current traces directly from protein sequence. **a**–**d**, Description of model signal generation. **a**, A protein sequence to be modeled. **b**, Calculation of the volume and charge, scaled, for all amino acids in the window of size 20. **c**, Parabolic weighting of the values within a window. **d**, Plotting the value *S* for each window, by computing the dot product of the parabolic weight array and the window array, to create the full model signal. **e**, Comparison between the nanopore signal of an example ionic current signal of PASTOR-TWAFH (black line) and the modeled signal generated for the same protein sequence (pink line). Model signal shown with the time axis aligned to the experimental trace using DTW. **f**, Distributions of the DTW distances between the real (experimental) signal traces and the model signals of the same sequence (pink), or between the real signal traces and the model signals of 10,000 random sequences derived from the same amino acid distribution as the real sequence (orange). n of experimental traces ranges from 27 to 55.

## Building an aminocaller for single-molecule sequencing

Sequencing PASTOR VRs would mark an important step in nanopore protein sequencing development. Moreover, sequencing synthetic protein constructs such as PASTORs could serve diverse technological applications, including protein barcoding^40,44^. We addressed this by initially training machine learning models to identify the single mutation present within a VR. This process consisted of filtering and scaling each of the raw signal traces, followed by segmentation of the VRs signal regions (**Fig. 4a**). To featurize the VR signals, we used a combination of manually curated features and DTW-distance features (**Methods**). The latter helped to quantify the pairwise similarity of all the VR reads in the training set. We next explored several classical and deep machine learning models and found that random forests most frequently achieved the highest accuracy. All classification analyses were then performed with a hyperparameter-tuned random forest evaluated on a fixed held-out test set, unless otherwise specified. We first started by evaluating amino acid discrimination across all pairs of amino acids (**Fig. 4b** and **Supplementary Fig. 10**). The highest accuracy classifications were achieved for pairs of amino acids with dissimilar volumes or when one was negatively charged, for example, tyrosine vs aspartic acid, which exhibited 100% discrimination accuracy. Some pairs of amino acids, such as leucine and isoleucine, proved to be more challenging due to their inherent physicochemical similarities. Amino acids with high variance signals, like cysteine, were also more difficult to distinguish from others. We then moved to training models to classify amongst particular sets of three amino acids (for example G, Y, and D), in which the model achieved 95% single-read accuracy (**Supplementary Table 1**). Expanding this to five-way classifications (for example G, V, W, R, and D), the model maintained high performance, achieving an accuracy rate of 86%. In the most challenging task of all 20-way amino acid classification, our top-performing model substantially outperformed a dummy classifier, obtaining an accuracy of around 28%, compared to the latter’s 5.5% (**Fig. 4c**). Lastly, when we considered top-N accuracy measurements, our model attained 67% accuracy for top-5 and 81% for top-8 accuracy in the 20-way classification task (**Fig. 4c**).

**Fig. 4.**
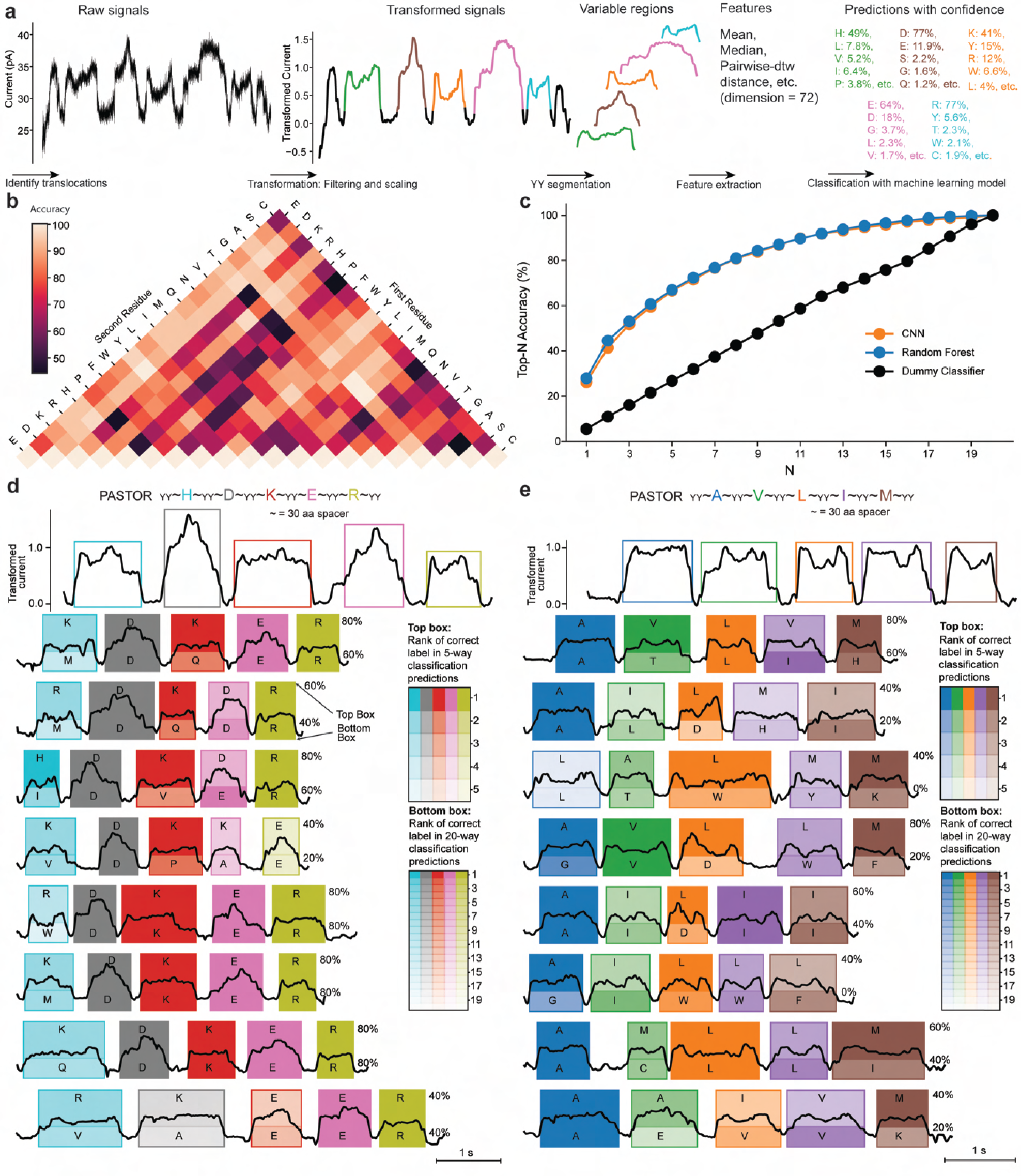
Single-molecule nanopore sequencing of single amino acid mutations. **a**, PASTOR VR classification pipeline. **b**, Heatmap showing test accuracies in discriminating between all pairs of amino acid VR mutations, averaged over five Random Forests. **c**, Accuracy in a 20-way classification when “accuracy” is defined as the correct label being in the top-N most probable classes. The dummy classifier chooses one label at random. Results averaged over 20 models. **d** and **e**, Example sequencing traces in the test set, for two PASTOR constructs HDKER and AVLIM. Transformed ionic current traces are plotted with a box around the variable regions defined by the segmenter. The color intensity of the boxes represents the ranking of the true class in the aminocaller’s prediction for each VR. For the 5-way classification task (top box shading), the classes are the 5 mutations found in that protein, while the 20-way classification task (bottom box shading) considers all possible amino acid classes. In each box, the letter corresponds to the model’s top prediction, with the top letter denoting the 5-way classification and the bottom letter indicating the 20-way classification. A darker shade implies a more accurate prediction, indicating that the correct label ranked high in the model’s predictions.

Building upon these results, we integrated our classifiers downstream of the PASTOR segmenter to develop an end-to-end PASTOR “aminocaller.” We then aminocalled a set of PASTOR reads from the classification test set (**Figs. 4d** and **4e**, and **Supplementary Fig. 11**). Overall sequencing accuracy per read averaged about 62% and 42% for the HDKER sequence, and roughly 51% and 21% for the AVLIM sequence, using five-way and 20-way classification models respectively. These results substantially exceed random baselines of 20% and 5%. These results expand the capabilities of nanopore protein reading, moving beyond discriminating among a limited set of amino acids to sequencing individual amino acids across a protein strand, albeit in a synthetic sequence background.

## Rereading single protein strands using an unfoldase slip sequence

Having established the aminocaller, we aimed to enhance the accuracy of our single-molecule sequencing approach by developing a method to reread single protein molecules. A multi-read strategy would facilitate the generation of consensus sequencing data at the individual molecule level. A previous study suggested that ClpX may have difficulty gripping particular polypeptide sequences, such as polyproline, on which the ClpXP complex showed slow degradation rates^33^. This prompted us to hypothesize that incorporating a “slippery” amino acid sequence near the N-terminal of a PASTOR would induce ClpX to momentarily lose hold of the strand (**Fig. 5a**). Consequently, the substrate protein would be free to rethread into the pore via electrophoresis. Rethreading would cease and enzyme-mediated translocation would resume once ClpX regains its grip on the substrate. To test this strategy, we constructed a new PASTOR (PASTOR-reread) with two important sequence features: 1) a proline-rich “slip” sequence repeat (EPPPP)_5_ positioned near the N-terminal, and 2) VRs separated by an increasing number of tyrosine residues, ranging from two to five (**Supplementary Fig. 1**). We reasoned that these distinctive tyrosine repeat lengths would aid in characterizing rereading events, given the distinct current levels produced by each repeat length. Reinforcing this hypothesis, our sequence-to-signal model predicted that this unique PASTOR-reread sequence design would generate a distinctive stair-step-like signal in the Y_n_-dips, enabling us to estimate the slip distance (**Supplementary Fig. 9**). Indeed, nanopore signals produced by PASTOR-reread generally exhibited repeated signal patterns that closely aligned with our model’s prediction before returning to open channel (**Fig. 5b** and **Supplementary Fig. 12**). By using the tyrosine repeat regions as a measure of slip length, we observed that slipping distances were most often either within short ranges (50–100 amino acids) or extending across the entire PASTOR unstructured region (>300 amino acids), accounting for roughly 40% and 30% of all rereads, respectively (**Fig. 5c**). The distribution of these slipping distances remained relatively unchanged regardless of ClpX concentration. However, the frequency of slips, whether partial or full, did vary with changes in ClpX concentration (**Figs. 5d** and **5e**). At a concentration of 1000 nM, we observed a median of 5.5 reads per PASTOR-reread molecule, whereas at a lower concentration of 8 nM, we observed 22.5 reads per PASTOR-reread molecule (**Fig. 5d**). These results suggest that higher ClpX concentrations may lead to multiple ClpX complexes loading onto a single PASTOR strand. This increased loading could potentially reduce the frequency of back-slipping, as downstream ClpX complexes might serve as a backstop for the upstream ClpX encountering the slip sequence region. While prior studies have documented ClpX back-slipping during unsuccessful protein unfolding^45,46^, these findings show direct evidence of purely sequence-dependent ClpX back-slipping. This suggests that this technique could also enable new single-molecule biophysical studies on protein-processive motors, paralleling nanopore research on DNA-processive motors^47,48^.

**Fig. 5.**
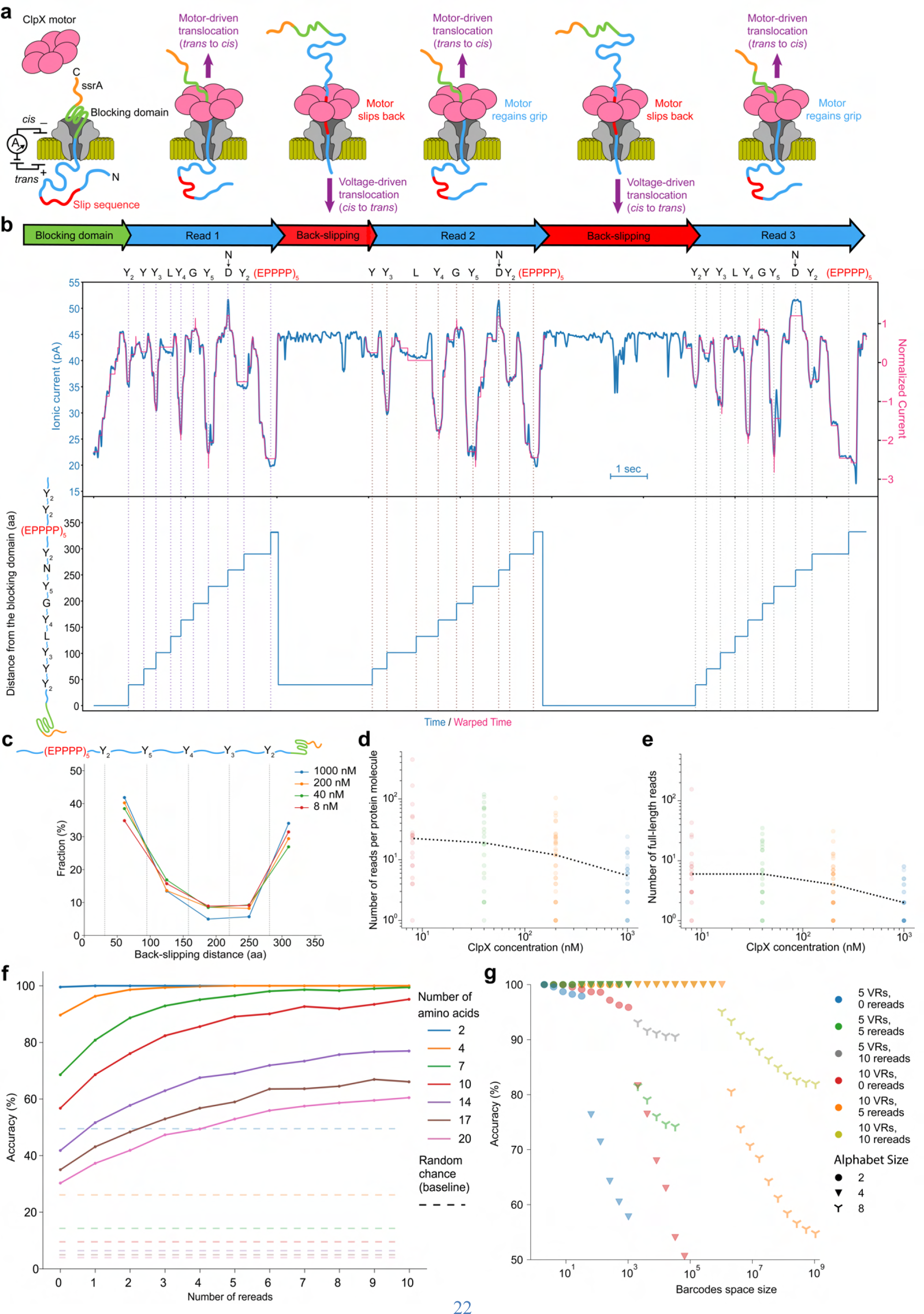
Rereading single protein molecules multiple times with an unfoldase slip sequence. **a**, Working model of rereading. The ClpX motor translocates PASTOR-reread to generate Read 1 (*trans* to *cis*). ClpX releases the protein strand upon reaching the slip sequence, and electrophoresis drives back-slipping (*cis* to *trans*). ClpX regains grip and ratchets the protein strand through the nanopore again. Recurrent back-slipping events produce subsequent rereads. **b**, Top box: Example trace of PASTOR-reread showing three near complete reread events (blue trace). Our model’s predicted signal for the PASTOR-reread sequence (pink trace) was aligned to each reread. The fourth VR contains an asparagine mutation, but the corresponding signal level consistently resembles aspartate in all three instances for this particular PASTOR-reread trace. The modeled sequence was changed to contain an aspartate to reflect the putative PTM. These rereading data provide additional evidence supporting the notion that the variability observed in asparagine VRs is attributed to post-translational deamidation of N to (iso)D, and not read-to-read variation. Bottom box: plot showing the approximate region of the strand that is within the nanopore over time. **c**, Estimated back-slipping distance for ClpX concentrations at 1000 nM (*n* = 141), 200 nM (*n* = 609), 40 nM (*n* = 777), and 8 nM (*n* = 999). The very first full-length read (Read 1) of each analyte protein molecule was excluded from this analysis. **d** and **e**, Number of all reads and full-length reads per PASTOR-reread molecule, respectively. The dotted lines indicate medians for ClpX concentrations at 1000 nM (*n* = 26), 200 nM (*n* = 37), 40 nM (*n* = 23), and 8 nM (*n* = 20). **f**, Simulated effect of rereading on 2 (Y, D), 4 (A, W, R, D), 7 (G, Q, W, F, R, D, E), 10 (A, G, V, N, Y, W, F, R, D, E), 14 (C, A, G, T, V, N, Q, M, Y, W, F, R, D, E), 17 (C, S, A, G, T, V, N, Q, M, I, Y, W, F, H, R, D, E), and 20-way (all 20 a.a.) classification tasks, compared to a baseline random classifier. Each value is the average over 100 train-test trials. **g**, Projected sequencing accuracy of barcode designs using the accuracies from f. Dots of the same color represent different amounts of bits allocated to error correcting codes (see **Methods**).

We next investigated the potential of single-molecule rereading in enhancing sequencing accuracy. While training our models similarly as before, we modified the testing approach. Instead of directly comparing the predicted classification to the actual label, we organized test VRs with the same amino acid into clusters of size n (representing n reads, or n − 1 rereads). Within each group, the class prediction with the highest average confidence was selected to represent the entire group. Using this approach, the accuracy for the all 20-way amino acid classification task improved from 28% to 61% (compared to a 5% random baseline) with 10 rereads (**Fig. 5f**). Likewise, the accuracy for a 7-way classification task improved from 66% to 99% (against a 14% random baseline). This method to estimating the effects of rereading offers promising indications of its potential to increase single-molecule sequencing accuracies. We also postulate that the use of reread data in practice could offer even greater accuracy improvements. This is especially likely in a scenario where all the reads are considered in summation by the aminocaller models, as opposed to merely leveraging individual prediction confidences.

## Projecting protein barcode space and decoding accuracy

Having established the capacity for high-accuracy sequencing through PASTOR rereading and the ability to design PASTOR proteins with customizable VR sequences, we simulated the PASTOR VR sequence space with varying constraints, with a view toward applications in protein barcoding. Based on the accuracy rates of our models (**Fig. 5f**), we computed the number of distinct barcodes that could be generated at a given accuracy level. This calculation considered varying degrees of rereading and two different VR segment numbers per protein barcode (5 and 10 VRs). A protein containing *L* VRs, where each VR could possess one of *a* possible amino acid mutations (where *a* = size of the mutation alphabet), would represent *L*∗log_2_(*a*) bits and could create a total of *a^L^* unique barcodes. However, by devoting some of those bits to linear error-correcting codes for parity checks, we can enhance the decoding accuracy, although it results in a reduced barcode space. For example, our findings indicate that with 10 VRs and 10 rereads, it’s feasible to generate libraries of over 1 million or 1 billion unique PASTOR barcodes that are decodable with a single-molecule accuracy >95% or >81%, respectively (**Fig. 5g** and **Supplementary Table 2**).

## Processive reading of folded protein domains

Progressing beyond synthetic, unstructured sequences, we next proceeded to evaluate the effectiveness of our *cis*-based unfoldase method on protein sequences that contain a folded domain. For this purpose, we analyzed a PASTOR protein with the titin I27_V15P_ domain, which consists of 89 amino acids arranged into 8 β-strands^49^, inserted into the third VR position (PASTOR-Titin; **Supplementary Fig. 1**). Unlike unstructured proteins, nanopore traces of PASTOR-Titin yielded an initial two-step electrophoretic nanopore capture state, indicating that the folded titin domain was first captured atop the nanopore at state ii and then electrically unfolded at transition state iii to produce the typical PASTOR capture signal at state iv manifested by the Smt3 domain atop the pore (**Figs. 6a** and **6b**). Following the addition of ClpX to the *cis* compartment, we observed a translocation signal corresponding to the leading VRs and YY regions, which are tethered to the C-terminus of the titin domain (state v). Subsequently, we observed a distinct and deep blockade state, which we interpreted as ClpX attempting to unfold the titin domain (state vi), which presumably refolds in the *trans* compartment after initial translocation. This deep blockade state often reverted back to the previous state, indicating an unsuccessful ClpX unfolding attempt and suggesting that ClpX slipped back on the protein strand^46,49^. Following a successful unfolding attempt, we observed putative translocation of the titin domain (state vii). After translocation of the titin domain, we observed characteristic signal features corresponding to the latter VRs and YY regions (state viii) before transitioning back to an open pore state (state ix) (**Fig. 6b** and **Supplementary Fig. 13**).

**Fig. 6.**
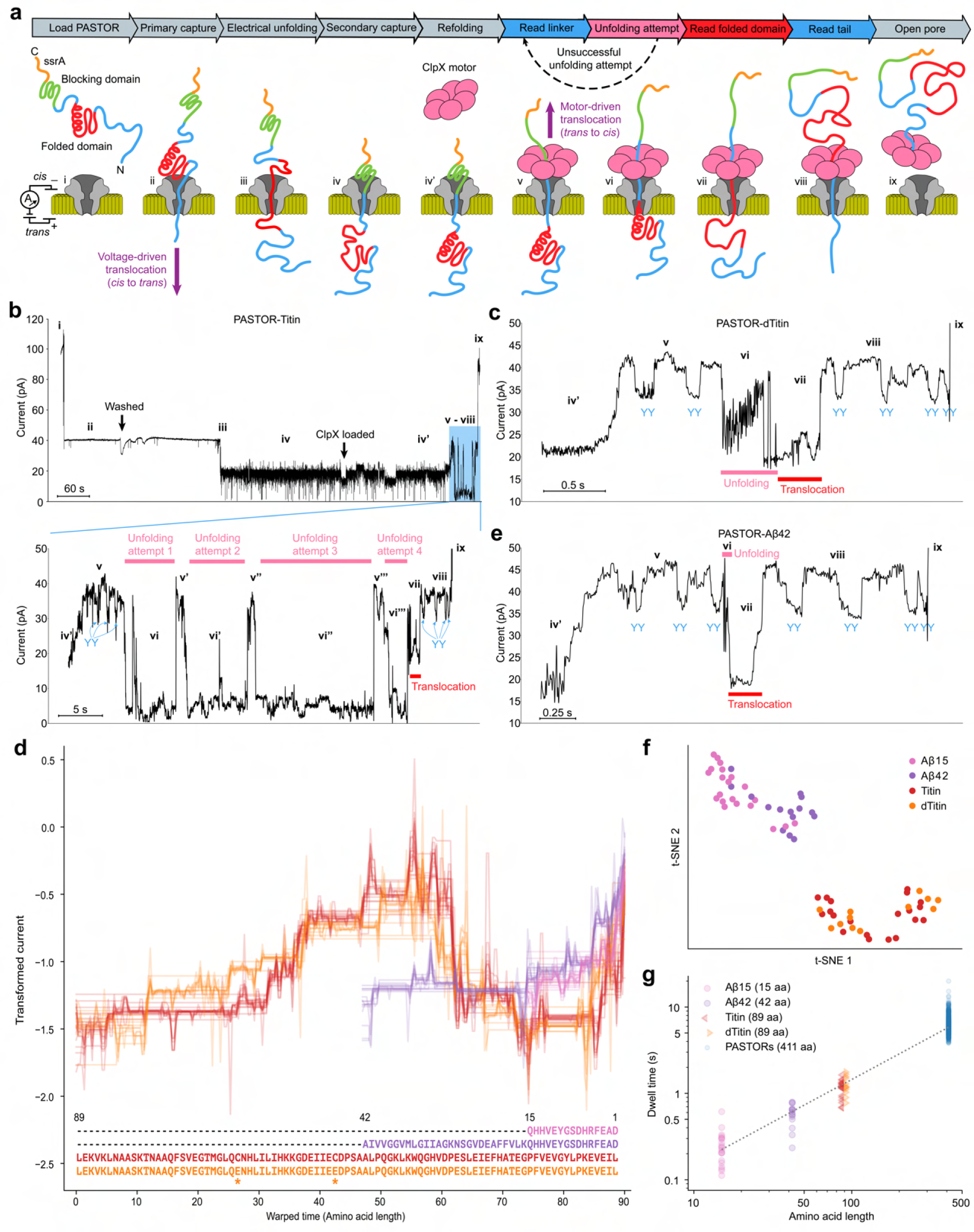
Processive reading of folded protein domains. **a**, Working model of ClpX-mediated processing of folded proteins. Assigned Roman numerals correspond to ionic current states in **b**, **c**, and **e**. **b**, Example trace of PASTOR-Titin. **c**, Example trace of PASTOR-dTitin. **d**, Ensemble traces of state vii of PASTOR-Aβ15 (pink, *n* = 21),-Aβ42 (purple, *n* = 15),-Titin (red, *n* = 20), and -dTitin (orange, *n* = 12). Protein sequences are shown in the C-to-N direction, and asterisks represent the C47E and C63E mutations between Titin and dTitin. **e**, Example trace of PASTOR-Aβ42. **f**, t-SNE plot based on pairwise DTW distances for state vii. **g**, Relationship between protein length and translocation time. State vii dwell time is plotted for Aβ15, Aβ42, Titin, and dTitin, as well as translocation time for the 8 PASTORs with no folded domain insert (*n* = 672). The dotted line was fitted with the mean dwell times of each protein class (slope corresponds to a translocation rate of 16 ms/aa or 63 aa/sec, R^2^=0.998).

To affirm our understanding of the unfolding and translocation states, we performed experiments using a variant of titin I27 (PASTOR-dTitin) with a destabilized tertiary structure, introduced through double-point mutations (C47E, C63E) on two buried cysteines^35,49^. Comparing PASTOR-dTitin with PASTOR-Titin allowed us to explore the effect of titin’s tertiary structure on the resulting current signals. PASTOR-dTitin generated traces that bore resemblance to those from PASTOR-Titin but with two notable differences (**Fig. 6c** and **Supplementary Fig. 13**). First, PASTOR-dTitin displayed unique signal features at the putative unfolding state vi, indicating structural disparities between the two variants. Second, states v–vi were observed only once in PASTOR-dTitin before the presumptive translocation state vii, in contrast to PASTOR-Titin where they were typically observed multiple times, leading to a substantial difference between the distribution of PASTOR-Titin and PASTOR-dTitin unfolding times (**Supplementary Fig. 14**). These differences can be attributed to PASTOR-Titin’s more stable, unfolding-resistant titin domain compared to that of PASTOR-dTitin. In PASTOR-Titin, repeated observations of states v and vi, which were not present in PASTOR-dTitin, support the conclusion that they result from unsuccessful unfolding attempts and ClpX back-slipping events, triggered by the stable titin domain. Moreover, despite their dissimilar structural stabilities, PASTOR-Titin and PASTOR-dTitin demonstrated similar signals during the putative translocation state vii (**Fig. 6d**). This similarity reflects their nearly identical primary amino acid sequences. The observation of similar signals at the proposed translocation state vii between PASTOR-Titin and PASTOR-dTitin, despite their differences in structural stability, underscores the role of the primary amino acid sequence in this process. It suggests that the primary sequence is the major determinant of the translocation signal through the nanopore, while structural variations exert a more significant influence on the preceding unfolding state.

We next tested PASTOR constructs with the amyloid beta protein 1–42 (PASTOR-Aβ42) and its shorter derivative 1–15 (PASTOR-Aβ15), which have distinct amino acid sequences and lengths from the titin domain. We reasoned that Aβ42 and Aβ15 would generate brief unfolding states as they are partially but not fully structured in their monomeric forms^50^. As expected, upon nanopore analysis they yielded ionic current traces similar to PASTOR-dTitin overall but with distinct features in unfolding state vi (**Fig. 6e** and **Supplementary Fig. 13**). Further, comparing their putative translocation states (state vii) using DTW distances, we observed that the signals generated by PASTOR-Aβ42 and PASTOR-Aβ15 share similarities, while being distinct from signals generated by PASTOR-Titin and PASTOR-dTitin (**Figs. 6d** and **6f**), reflective of the translocation state being dependent on protein primary sequence. Overall, the dwell times of these different states also corresponded well with their respective sequence lengths across all the PASTOR proteins, suggesting a translocation rate of ∼63 amino acids/second (average dwell time of ∼16 milliseconds/amino acid) (**Fig. 6g**). This is close to previous estimates of ClpX translocation speed, and the observation that the rate of ClpX-mediated protein translocation is relatively constant regardless of protein sequence^35,51^. Altogether, these results provide evidence that the unfolding state vi and translocation state vii independently record structural and sequence information, respectively, showing the potential of this technique to enable multimodal protein analysis on a single platform.

Finally, we assessed our predictive model using these proteins. As the current model does not factor in the signal features linked with unfolding, we analyzed the signal segment following the unfolding state until the completion of the translocation (i.e. states vii–viii). Using the same comparison technique as previously implemented for the PASTOR protein models, we found that the average *DTWDS* for the PASTORs containing folded domains ranked in the top 0.04% of the best matches within *R* (**Supplementary Fig. 15**).

## Discussion

We have introduced a new approach for single-molecule reading of long protein strands using nanopores and an unfoldase motor protein. This method achieves single-amino acid sensitivity and demonstrates the capability to sequence amino acid substitutions across long protein strands. This could immediately open up new opportunities for the advancement of protein barcoding technology, as we project the ability to design large libraries of synthetic peptide sequences (>1B) that can be decoded with high accuracy. Further, the unfoldase enables processive translocation of folded protein domains for full-length analysis, an important result as we move towards reading of natural protein molecules. Although, it’s possible that some protein domains may exhibit greater resistance to unfolding than the small set explored in this study, necessitating additional strategies for their analysis, such as the use of denaturants^52^.

Stemming from this method, we have laid the groundwork for a biophysical model capable of simulating nanopore signals that are generated as individual protein sequences are pulled through the nanopore by the unfoldase. This result could eventually enable a “lookup table” approach reminiscent of mass spectrometry, facilitating full-length, single-molecule protein identification and fingerprinting. This strategy would entail creating a database comprising simulated nanopore signals associated with various target protein sequences, potentially encompassing an entire proteome, and could be generated computationally given a robust sequence-to-signal predictor. The raw data derived from actual nanopore readings could then be compared against this database to identify the best protein match based on raw signal alignment, for example by utilizing DTW. This offers an attractive alternative to sequence-based matching, which is a complex task due to the challenge of accurately mapping ionic current to specific amino acid sequences, given the large number of amino acids contributing to the nanopore signal at any one time. However, we note that the CsgG pore variant (R9.4.1) employed in our study was optimized for DNA sequencing^37^. Given the substantial physicochemical differences between nucleic acids and peptides, future efforts will likely benefit from the exploration of new pore variants specifically optimized for protein sequencing.

This method has additional challenges that can also be addressed in future work. The two-step flow cell loading process currently limits the throughput of data collection. We envision an improved system: one that operates continuously, where the unfoldase is prebound to the protein analyte yet blocked from initiating its unfolding activity until the protein strand is captured in the pore. This approach would be similar to the strategy devised for nanopore sequencing of DNA^53^ and promises to enhance throughput. Moreover, controlling the electroosmotic flow to stretch neutral and positively charged protein regions could increase and standardize strand persistence length within the pore^54^. This adjustment could make the method more sensitive to primary sequence and enhance the correlation between the observed and model-predicted scores. This becomes especially important when modeling a broader range of sequence motifs that encompass secondary structures, such as alpha helices, a factor our current model does not account for. Likewise, adding the ability to model signal features linked to the unfolding of tertiary structures would increase the utility of our modeling approach. Finally, as we begin to turn our attention to natural proteins, this methodology will require synthetic N- and C-terminal sequences, which can be appended using existing termini-specific chemical conjugation techniques^31,55,56^. This requirement mirrors the adapter ligation step that is required for most DNA sequencing methods. In conclusion, this work serves as a stepping stone towards full-length protein identification, capable of achieving the highest level of proteoform resolution. Additionally, it promises immediate advancements, particularly in the context of protein barcoding applications.

## Acknowledgments

We thank Melissa Queen for advice on the barcode error correcting code, and additional members of the Molecular Information Systems Lab for feedback on this work. We also thank James Graham, Kiran Sabharwal, Jayne Wallace, Richard Gutierrez and others at Oxford Nanopore Technologies for helpful discussions and providing the configurable MinION runscript.

## Funding

National Institutes of Health grant R01HG012545 (JN)

National Science Foundation grant EF-2021552 (JN)

Oxford Nanopore Technologies sponsored research agreement AM03 (JN)

Japan Society for the Promotion of Science Overseas Research Fellowships (KM)

AMED Scholarship Program for Young Researchers related to Drug Discovery and Development (KM)

Marilyn Fries Endowed Regental Fellowship in Computer Science & Engineering (DKH)

## Author contributions

KM, JW, KK, GR, YF, and NC performed wetlab experiments. DKH developed the data processing pipelines. DKH and SY developed computational models. KM, DKH, JW, and KK performed data analysis. JN supervised the project. JN and KM conceived the project. KM, DKH, and JN wrote and edited the manuscript.

## Competing interests

Provisional patents covering aspects of this work have been filed by the University of Washington. JN is a consultant to Oxford Nanopore Technologies. The remaining authors declare no competing interests.

## Data and materials availability

Protein expression plasmids will be made available in Addgene. Data can be found on https://github.com/uwmisl/PASTOR-sequencing. Custom MinION MinKNOW run scripts can be obtained from Oxford Nanopore Technologies upon request.

## Code availability

Code for analyses is available at https://github.com/uwmisl/PASTOR-sequencing.

## Methods

### Expression and purification of proteins

Plasmids for analyte proteins were constructed with gBlocks (Integrated DNA Technologies) inserted into the pET-49b(+) plasmid (Novagen), with a dihydrofolate reductase domain, a polyhistidine tag, and a TEV cleavage site upstream of the sequence encoding an analyte protein. The NEBuilder HiFi DNA Assembly and Q5 site-directed mutagenesis kits (New England Biolabs) were used for plasmid construction. Cloning was performed using NEB 5-alpha competent *E. coli* cells. Plasmid sequences were verified by Sanger sequencing through Genewiz. Protein expression was induced overnight at 30°C with BL21 (DE3) *E. coli* cells in Overnight Express Instant TB medium (Novagen). Proteins were purified via immobilized metal affinity chromatography (IMAC) with TALON metal affinity cobalt resin and its associated buffer set (Takara), following the manufacturer’s instructions. Proteins were cleaved with TEV protease (New England Biolabs) and further purified via reverse IMAC. Purified proteins were concentrated using ultracentrifugal filters with a 10 kDa cutoff (Amicon) and stored over the short term at 4°C or over the long term at −80°C until use. For asparagine deamidation buffer conditions, analyte proteins were incubated overnight in 100 mM sodium bicarbonate buffer (pH 9.6) at 25°C to catalyze deamidation. A covalently linked hexamer of an N-terminal truncated ClpX variant (ClpX-ΔN_6_)^57^ was prepared using the BLR *E. coli* strain as described previously^35^. In brief, cells were grown to an A_600_ of ∼0.6 in LB medium and then incubated in the presence of 0.5 mM isopropyl β-D-1-thiogalactopyranoside at 23°C for ∼3 hr to induce ClpX expression. ClpX was purified via IMAC and anion exchange chromatography. Purified ClpX was stored at −80°C in small aliquots until use.

### MinION experiments

All experiments were performed on the MinION platform using R9.4.1 flow cells. Run conditions were set with a custom MinKNOW script (available from Oxford Nanopore Technologies) at a temperature of 30°C and a constant voltage of −140 mV with a 3 kHz sampling frequency, except for initial proteins P1–P4, in which runs were performed at a constant voltage of −180 mV with a 10 kHz sampling frequency. Using the priming port, flow cells were first washed with 1 mL *cis* running buffer (200 mM KCl, 5 mM MgCl2, 10% glycerol, and 25 mM HEPES-KOH, pH 7.6) and then loaded with 200 uL of protein analyte (final concentration: ∼0.5 µM protein) in *cis* running buffer. Following observation of protein captures in the pores, flow cells were washed with 1 mL of *cis* running buffer to remove uncaptured proteins and subsequently loaded with 75 uL of *cis* running buffer supplemented with 4 mM ATP and 200 nM of ClpX-ΔN_6_ unless otherwise specified.

### Nanopore signal analysis

#### Preprocessing

To assist in identifying ClpX-mediated protein translocations, we established detection thresholds using specific statistical parameters (standard deviation, median value, standard deviation of the mean of windows, and the ratio of values relative to the open pore value) indicative of translocation to ionic current blockades preceding a return to open channel. This analysis was used to assist the process of manually checking traces for translocations, and translocations with substantially high noise or disruptions were discarded. PASTOR proteins were auto-segmented as described below, with the exception of PASTOR proteins containing folded domains and PASTOR-reread, which were manually segmented. PASTOR-reread rereads with a complete Y_2_-Y_3_-Y_4_-Y_5_-Y_2_ signal were assumed to be full-length reads with a back-slipping distance of 310 aa. Partial rereads missing the signal(s) of the C-terminal Y_2_, Y_3_, Y_4_, and Y_5_ were assigned to back-slipping distances of 250, 188, 125, and 61, respectively. All figures with raw traces (those shown in pA) had a low-pass bessel filter applied using SciPy with N = 10 and W_n_ = 0.025. Before use in data analysis, traces were transformed (as in **Figs. 1c–d**, **2c–e, 2g, 4, 5f–g, 6d, 6f)** by applying a low-pass bessel filter with N = 10 and W_n_ = 0.03, except for PASTOR-Titin, -dTitin, -Aβ42, and -Aβ15 with W_n_ = 0.25, average downsampling by a factor of 50 for proteins P1–4, 20 for the 8 PASTORs, and 10 for the other proteins, and then scaling. To scale, the segment was split into tenths, and the median of the minimums of each tenth and the median of the maximums of each tenth were used as the min and max, respectively, to perform min-max scaling. Unless otherwise specified, ‘normalized’ refers to z-score normalization, as in ‘normalized current’ when comparing a model signal to experimental signals.

#### Signal alignment

To align signals, we used DTW^58^, and normalized DTW distances by dividing by the sum of the lengths of the two signals. To describe the similarity of a set of traces, we computed the DTW distance between all pairs of traces. In t-SNE plots, we then clustered traces on the vector of its DTW distances to all other traces. To create consensus signals, we used Dynamic Time Warping Barycenter Averaging^39^, with medoids selected to minimize the sum of squares of DTW distances to other signals. **Figs. 1c** and **6d** show all traces aligned to the consensus signal with warped time scales adjusted by protein length.

#### Protein sequence-to-signal model

To describe the amino acids, we used their volumes^59^ and their charge at pH = 7.6, where the histidine residue is assumed to be neutral. The model signal of a sequence [aa_1_, aa_2_,…, aa_n_] is calculated by computing the signal, S_i_, for each of the *n–*19 windows of width 20. The vector **X*_i_*** describes the window starting at index *i* in the sequence. The *j*^th^ index in **X*_i_*** is V_c_∗Volume(*aa_i+j_*) + C_c_∗Charge(*aa_i+j_*), for 0 ≤ *j* < 20. To weight the values in **X*_i_***, we use a vector PW of length 20 containing values representing a negative, centrally positioned parabolic curve. S_i_ is then computed as the dot product of **X_i_** and **PW**. The weights between charge and volume, V_c_=4∗10^-4^ and C_c_=1.2∗10^-4^, were determined empirically to minimize the DTWDS of a training subset of protein traces.

#### YY-Segmentation

To identify the YY dips and VRs, a single PASTOR trace was manually segmented into each colored section shown in **Fig. 2a**, and the remainder of the traces were aligned to it with DTW. The corresponding regions were assigned the label from the one manually segmented trace (**Supplementary Fig. 5**).

#### VR classification

We used scikit-learn to develop and test classical machine learning models and Pytorch to develop and test convolutional neural network models. The test set was composed of all current traces from a given set of experiments to best approximate an out-of-sample test set. The experiments were selected using linear programming (Python package Pulp) to ensure at least 12 VRs with each amino acid in the test set, while minimizing the test set size. The number 12 was selected, because it gave the closest to an 80-20 train-test split. 79.6% of the VRs were in the training set, and 20.4% were in the testing set. In classification tasks where only VRs corresponding to a subset of amino acids were used, the test set is composed of a subset of this test set. We performed hyperparameter tuning with scikit-optimize on the training set using 5-fold cross validation. The optimal parameters were n_estimators=250, min_samples_leaf=2, max_features=’log2’, max_depth=20, ccp_alpha=0.0001, class_weight=’balanced_subsample’, and criterion=’gini’. All results from **Figs. 4b-e** are from models evaluated on the test set. The results from **Figs. 5f** and **5g** are using random forest without hyperparameter tuning, and are the results from 100 randomly selected 80-20 train-train splits. This was necessary to sufficiently estimate the accuracy with a large number of rereads, given the data limitation and the need to group samples in the test set. All VRs containing an asparagine with a maximum transformed value above 1.3 had their labels changed to aspartate. In training all classical models, we upsampled minority classes, such that there was an equal representation of all classes in the training set. When training the CNN in **Fig. 2c**, we weighed the loss inverse proportion of the label’s class’s representation in the training set. To featurize the VRs, we performed PCA on the vector of its DTW-distances to all VRs in the training set, to reduce the size of the vector to 64. We also used the median, max, middle, mean, dip, mean absolute value of the derivative, and median absolute value of the derivative of the transformed signals, in addition to the standard deviation of the raw (unfiltered, unscaled) signal. The CNN had the transformed signal as input. It was trained with a stochastic gradient descent optimizer with a learning rate of 0.01, had four convolutional layers followed by a GRU and then a fully connected layer, and was initialized with Kaiming initialization. Max pooling and a ReLU activation function were applied after each convolutional layer. The dummy classifier was implemented with scikit-learn’s dummy classifier with default parameters.

#### Barcode error correction

To calculate the accuracy of barcode identification when using linear error correcting codes, we started with our accuracy, *p_VR_*, of identifying a VR given an alphabet size, *a*, of 2, 4, 8, and 16. For a given *a* and number of VRs, *L*, we calculated the number of bits *n*= *L*∗log_2_(*a*) that could be encoded in a protein. We simulated the accuracy with error correction, *p’*, when *n-k* of the bits were allocated to linear error correcting codes, for all integers *k*=1 to *n*. We did this by conducting 50,000 trials of 1) encoding a random integer from 0 to 2^k^ with a generating matrix into a message of *n* bits; 2) randomly, independently, with probability *p_VR_*, changing each of the 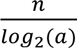 consecutive sets of log_2_*(a)* bits in the encoded message (to a different set of bits of the same length) to simulate misclassifying one VR; and 3) decoding the number with syndrome decoding. We calculated *p*’ to be the percent of trials where the decoded number equaled the original random number.

## Statistical Information

| Residue | A | C | D | E | F | G | H | I | K | L | M | N | P | Q | R | S | T | V | W | Y |
| --- | --- | --- | --- | --- | --- | --- | --- | --- | --- | --- | --- | --- | --- | --- | --- | --- | --- | --- | --- | --- |
| Total | 82 | 63 | 69 | 79 | 56 | 81 | 63 | 89 | 70 | 89 | 98 | 81 | 72 | 91 | 79 | 82 | 75 | 88 | 56 | 62 |
| Training set | 65 | 51 | 52 | 62 | 44 | 65 | 46 | 72 | 53 | 72 | 81 | 65 | 60 | 75 | 62 | 66 | 59 | 71 | 44 | 50 |
| Testing set | 17 | 12 | 17 | 17 | 12 | 16 | 17 | 17 | 17 | 17 | 17 | 16 | 12 | 16 | 17 | 16 | 16 | 17 | 12 | 12 |

Counts of PASTOR current traces:
| HDKER | GNQST | FYWCP | AVLIM | VGDNY | TWAFH | PRMQE | KSILC |
| --- | --- | --- | --- | --- | --- | --- | --- |
| 36 | 48 | 29 | 55 | 33 | 27 | 43 | 34 |

## Supplementary Information

Supplementary Figs. 1–15

Supplementary Tables 1 and 2

---

**Supplementary Fig. 1.**
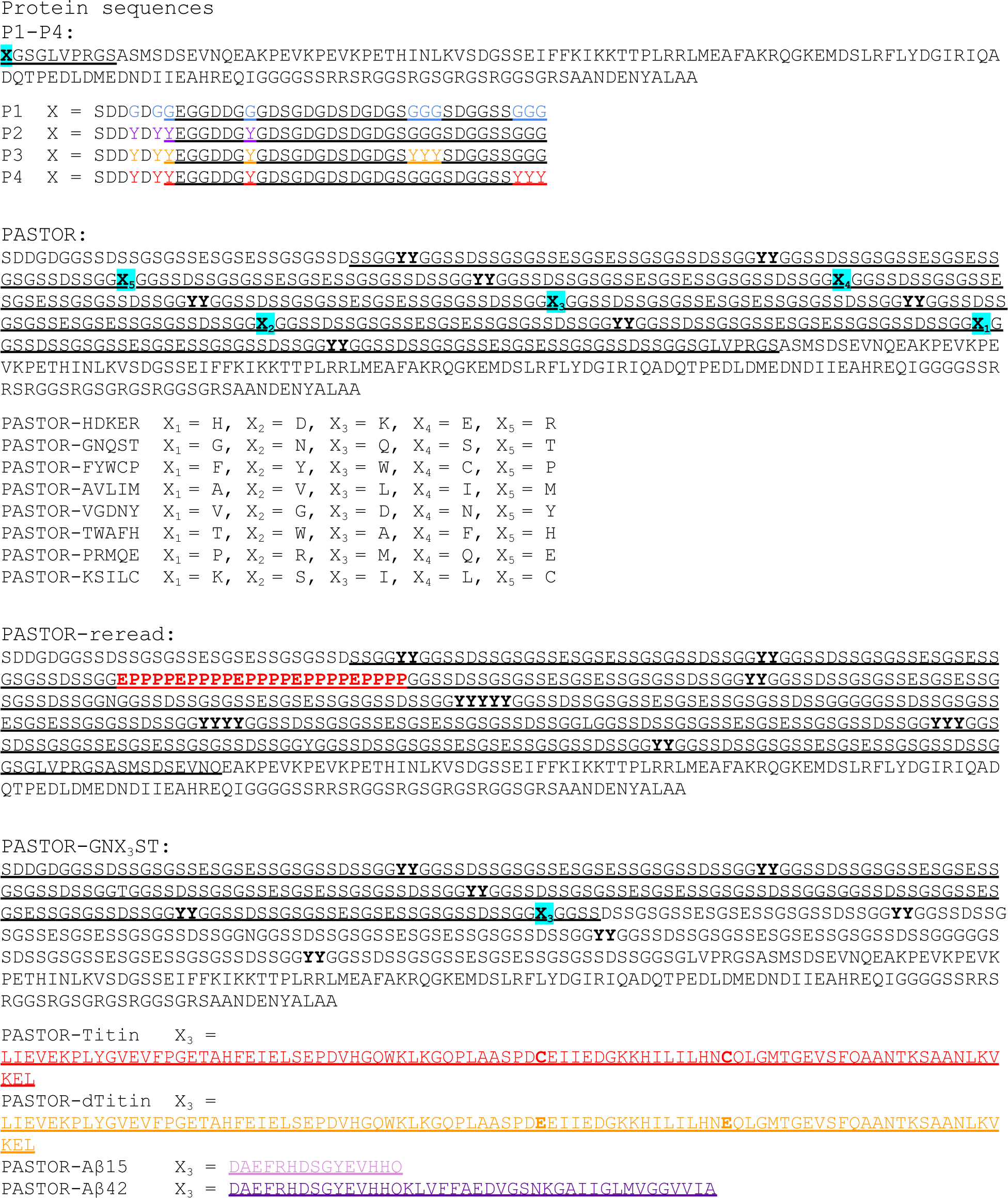
Protein sequences used in this study. Protein sequences are shown in the N-to-C order, which is opposite of the direction of ClpX-mediated translocation. The underlined residues represent the amino acid sequence used when modeling. Variable regions are highlighted in cyan.

**Supplementary Fig. 2.**
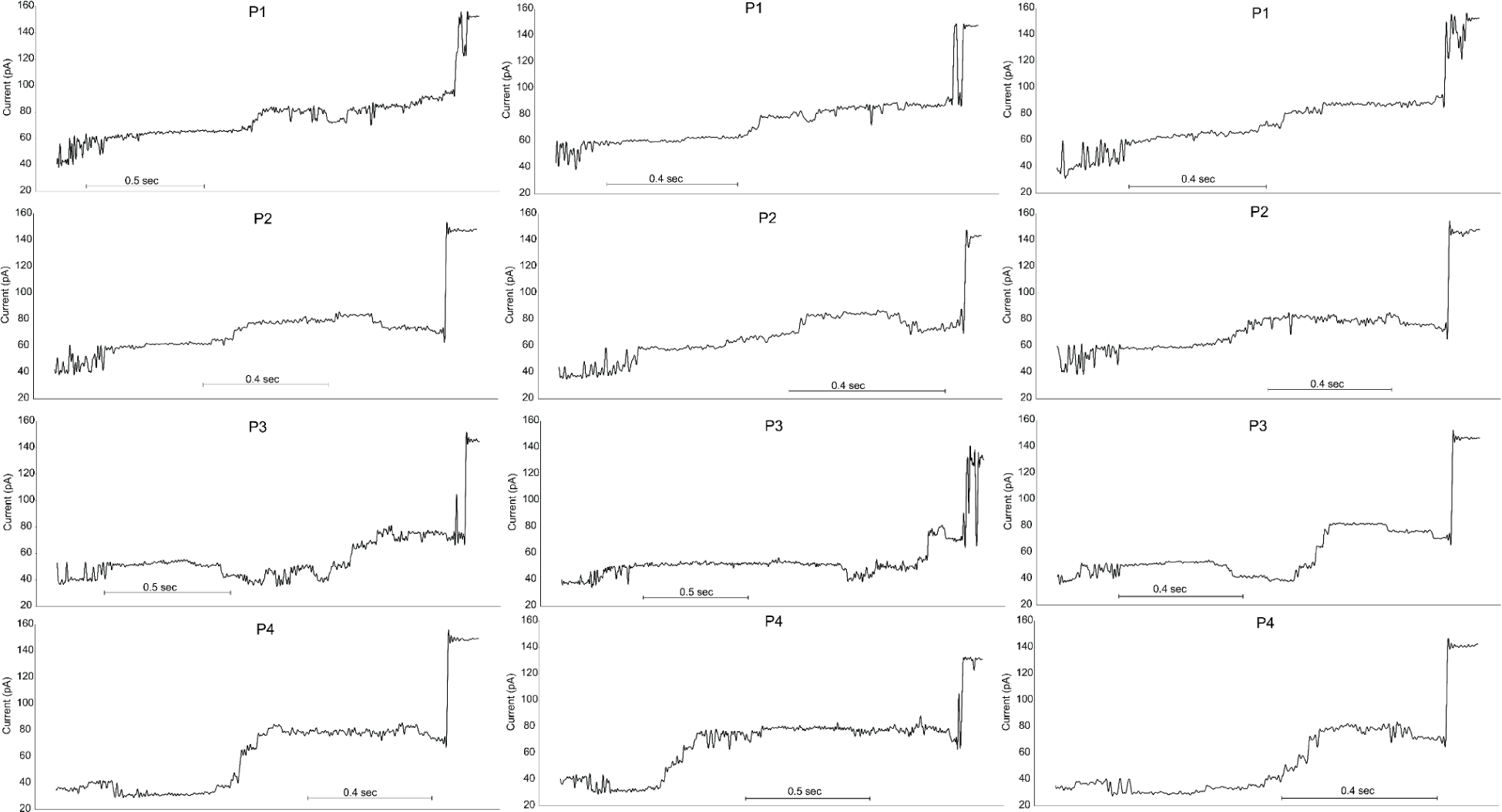
Example filtered traces of proteins P1 (top row), P2 (second row), P3 (third row), and P4 (fourth row).

**Supplementary Fig. 3.**
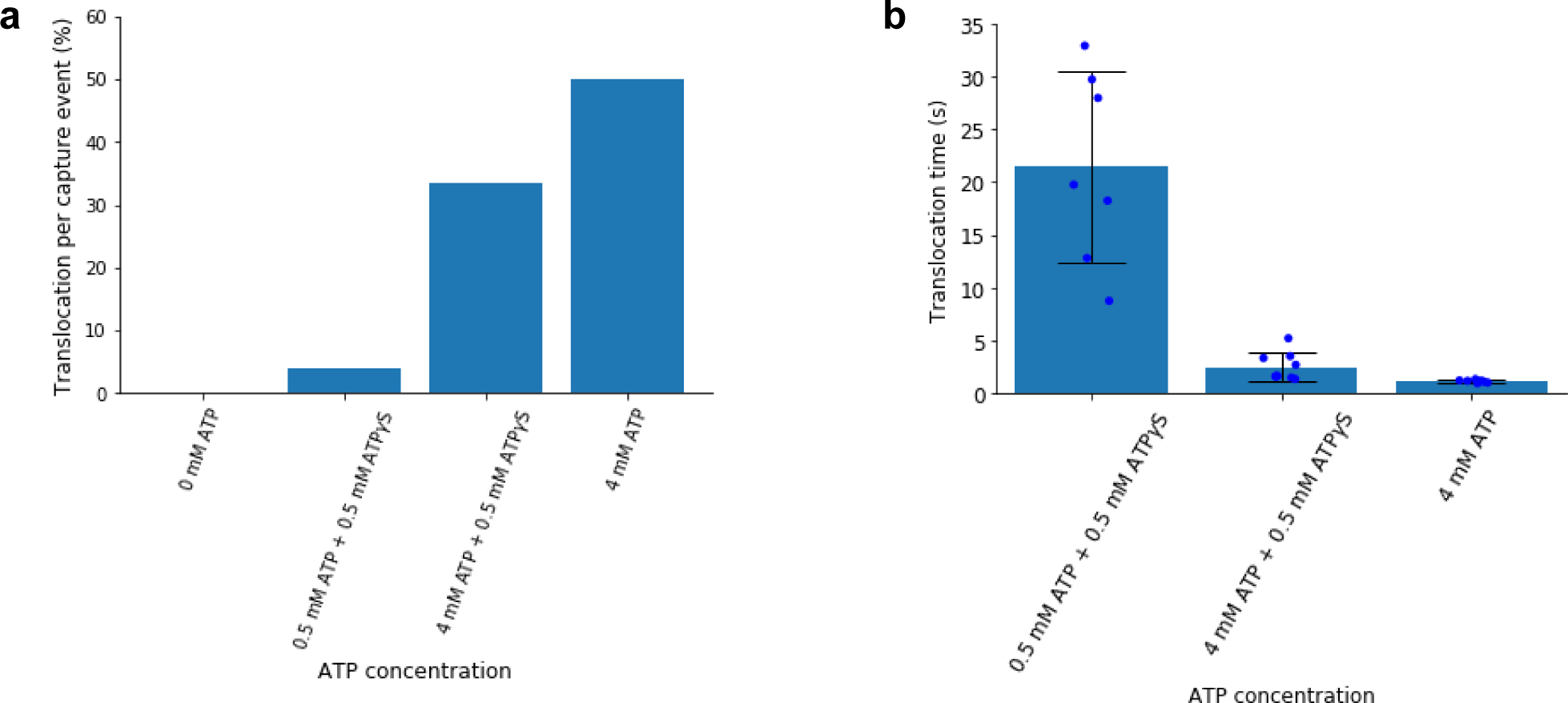
ClpX-mediated translocation is ATP-dependent. **a**, Fraction of ClpX-mediated translocation events observed following capture events in the presence of no ATP (*n* = 230), 0.5 mM ATP + 0.5 mM ATPγS (*n* = 180), 4 mM ATP + 0.5 mM ATPγS (*n* = 27), or, 4 mM ATP (*n* = 16). **b**, ClpX-mediated translocation time in the presence of 0.5 mM ATP + 0.5 mM ATPγS (*n* = 7), 4 mM ATP + 0.5 mM ATPγS (*n* = 9), or 4 mM ATP (*n* = 8). Error bars denote standard deviations.

**Supplementary Fig. 4.**
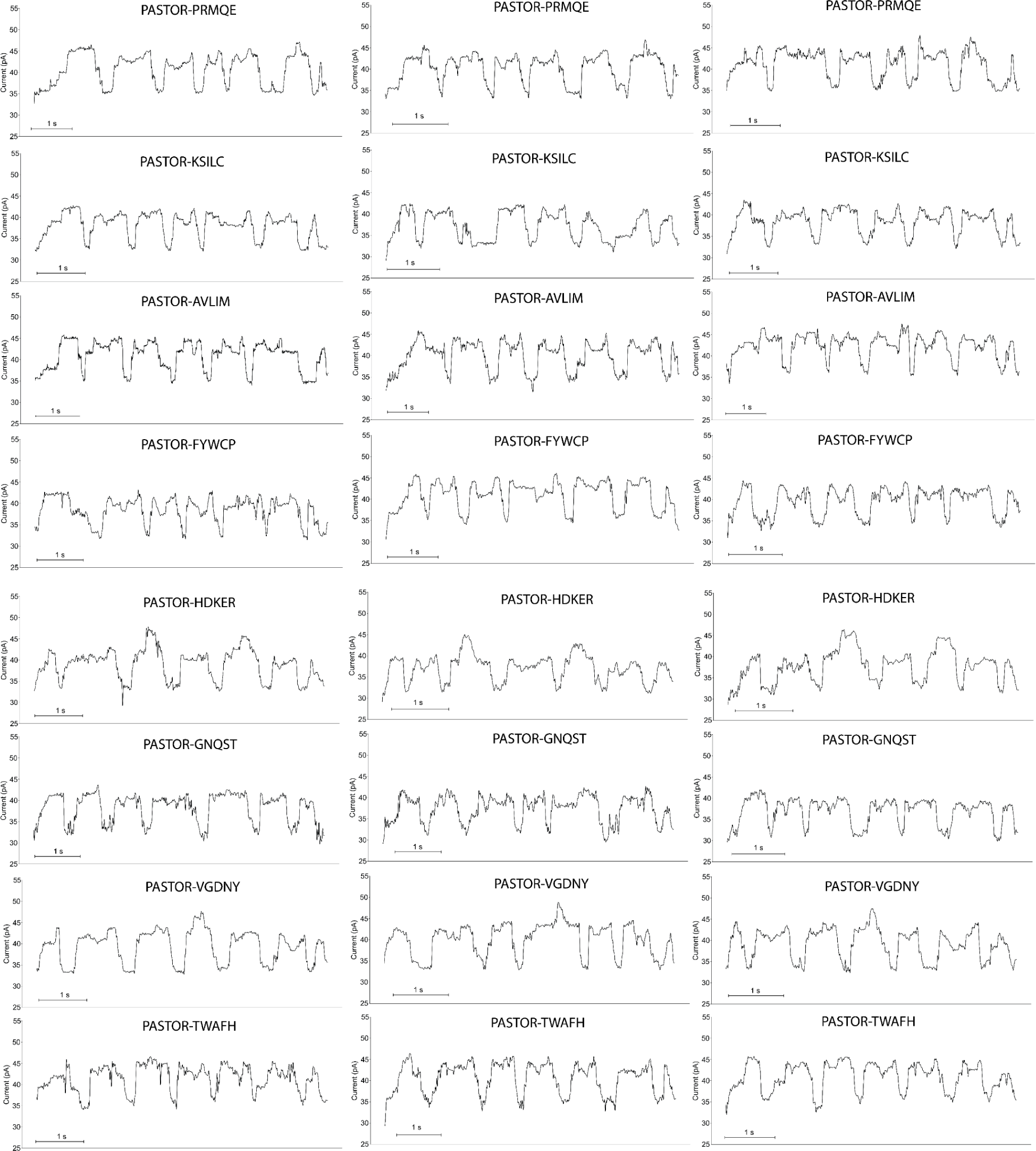
Example filtered traces of PASTORs.

**Supplementary Fig. 5.**
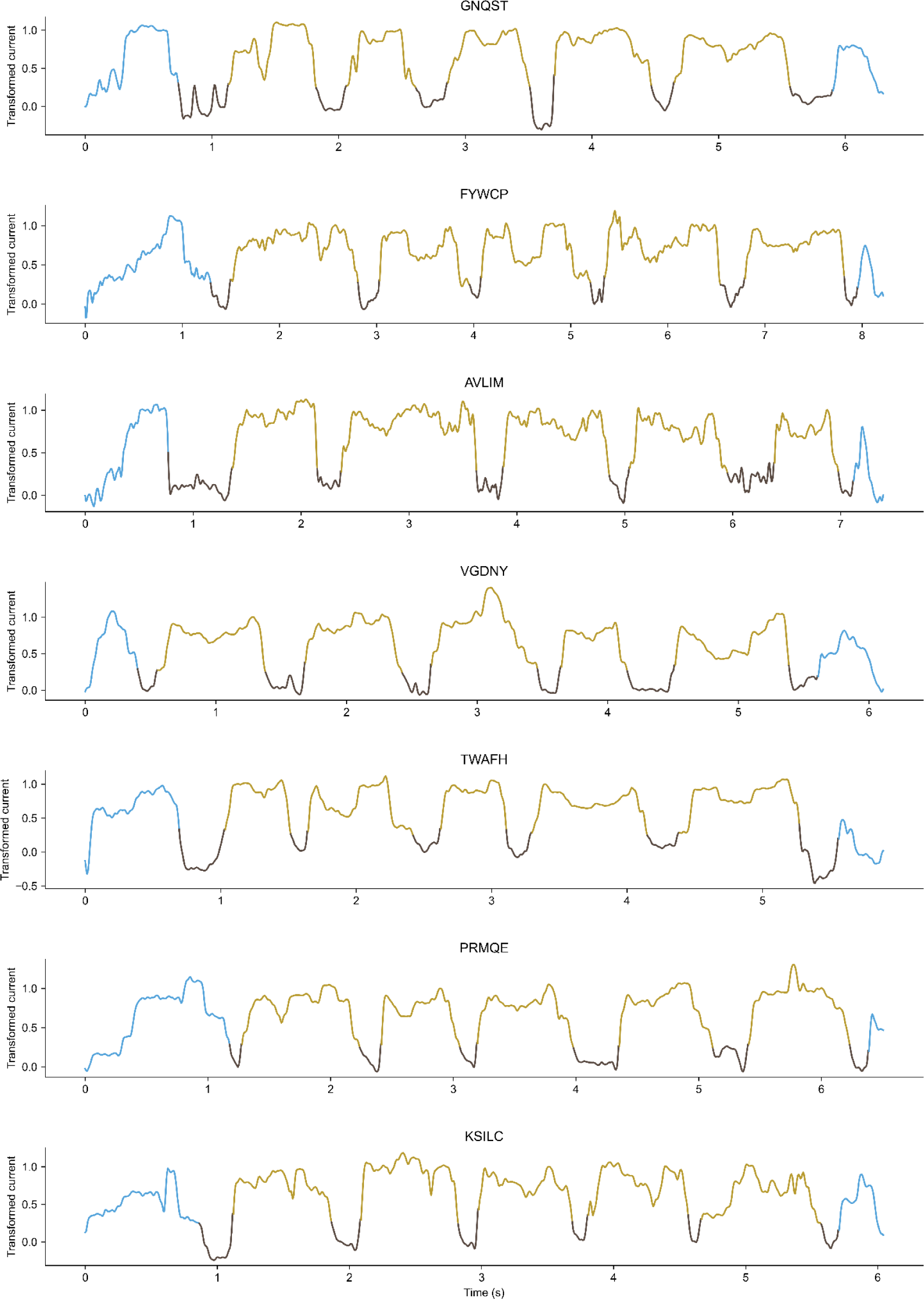
YY-Segmented traces of representative PASTOR ionic current signals. Dark brown shows YY dips and tan shows VRs.

**Supplementary Fig. 6.**
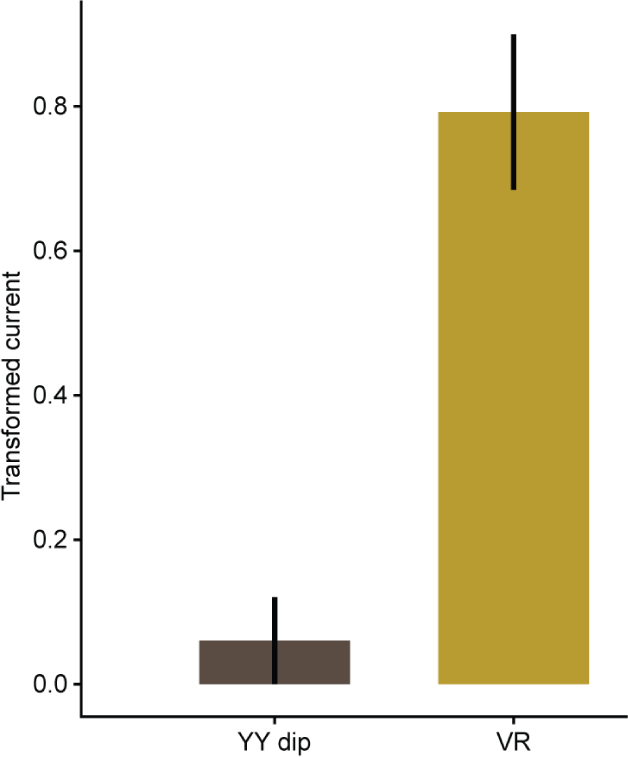
Mean transformed current levels of YY and VR PASTOR segments. Error bars denote standard deviation. n=1828 for YY dips and 1525 for VRs. There were a total of 305 PASTOR traces analyzed.

**Supplementary Fig. 7.**
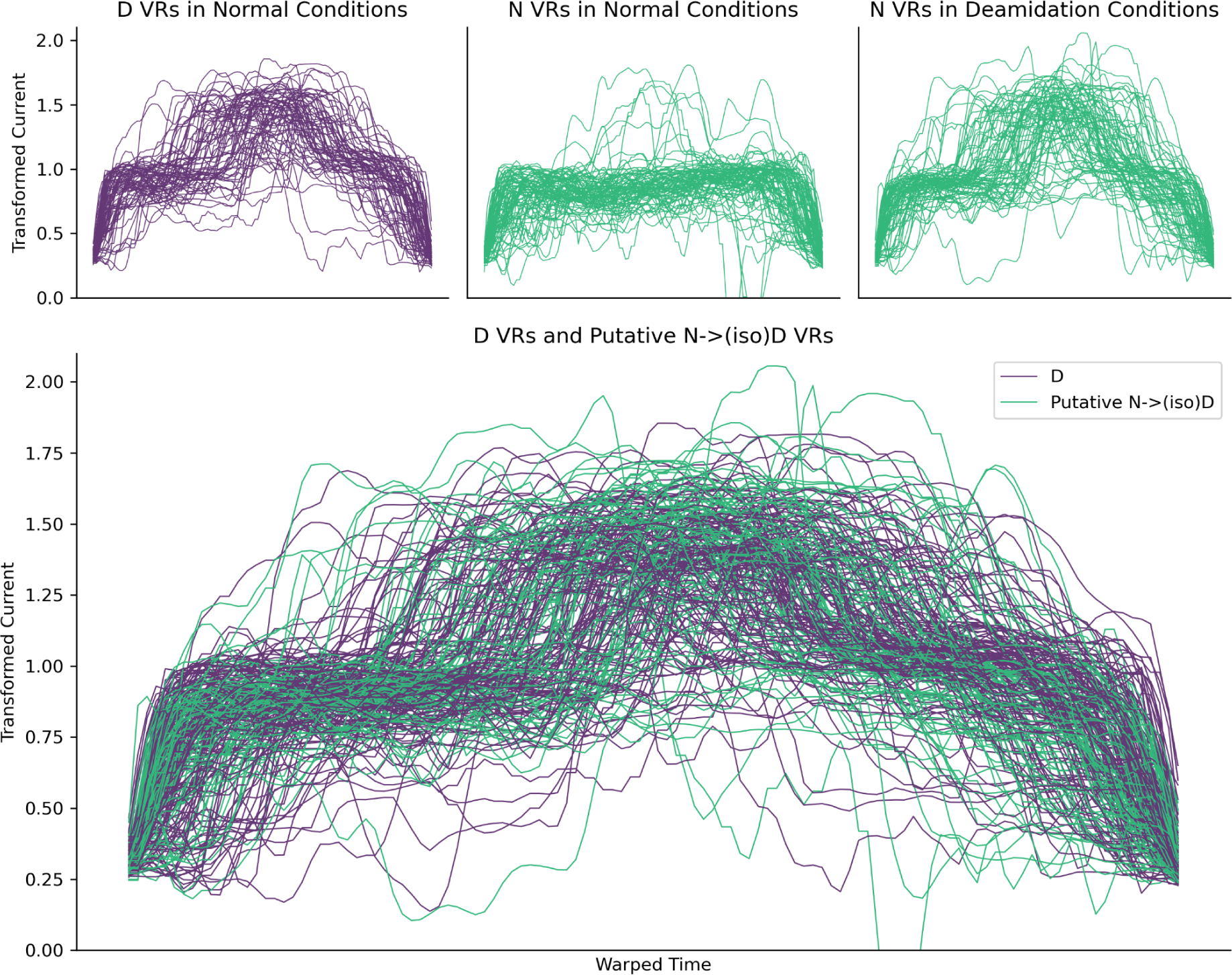
Transformed asparagine (N) and aspartate (D) VR signals in normal and deamidation catalyzing conditions. Putative modified asparagine VRs are those with a max transformed current value >1.3.

**Supplementary Fig. 8.**
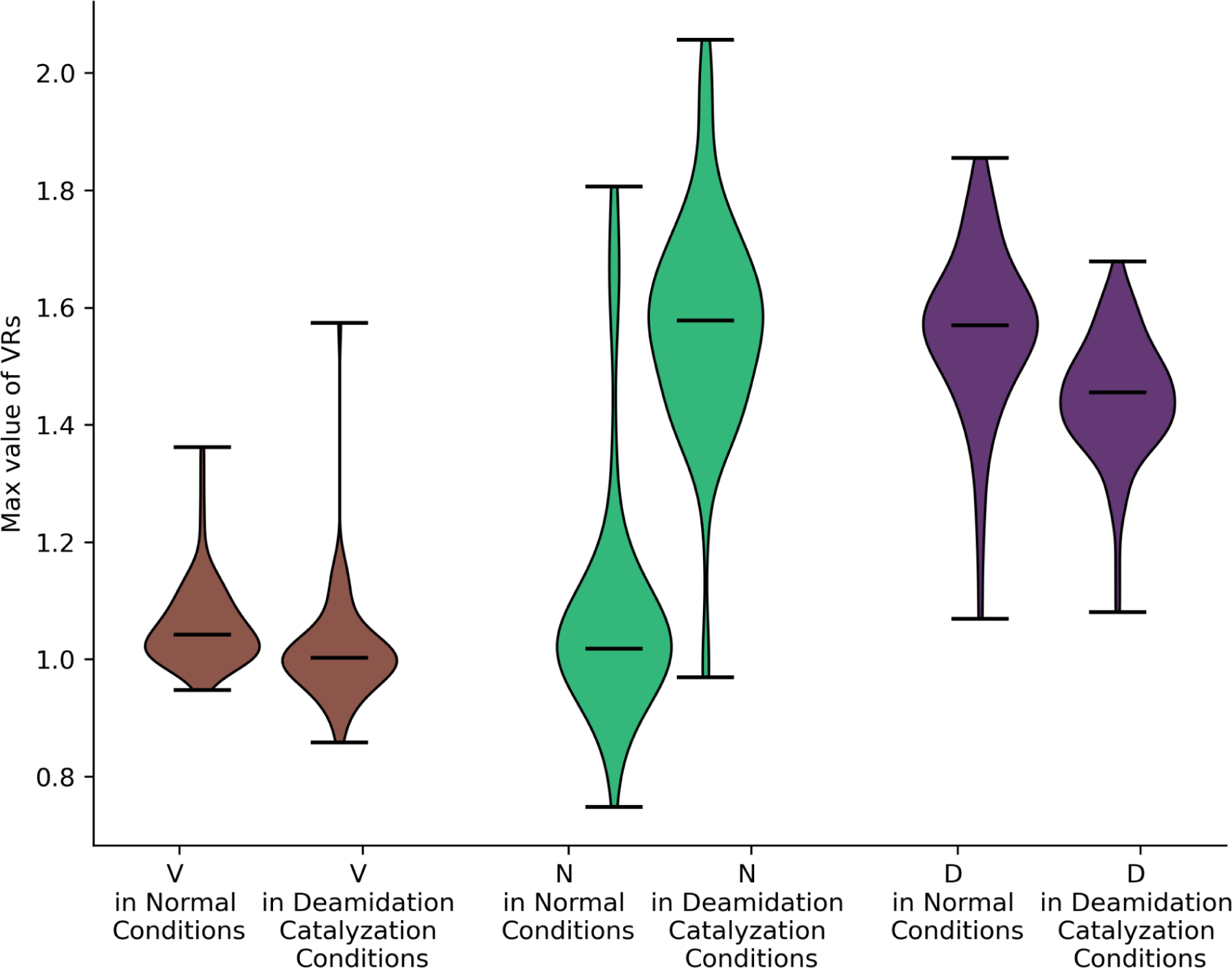
Violin plots showing distribution of the maximum height of transformed VRs, in normal and deamidation catalyzing conditions, for valine (V, brown), asparagine (N, green), and aspartate (D, purple). Horizontal lines denote min, median, and max. *n* = 88, 77, 81, 77, 68, and 77 from left to right.

**Supplementary Fig. 9.**
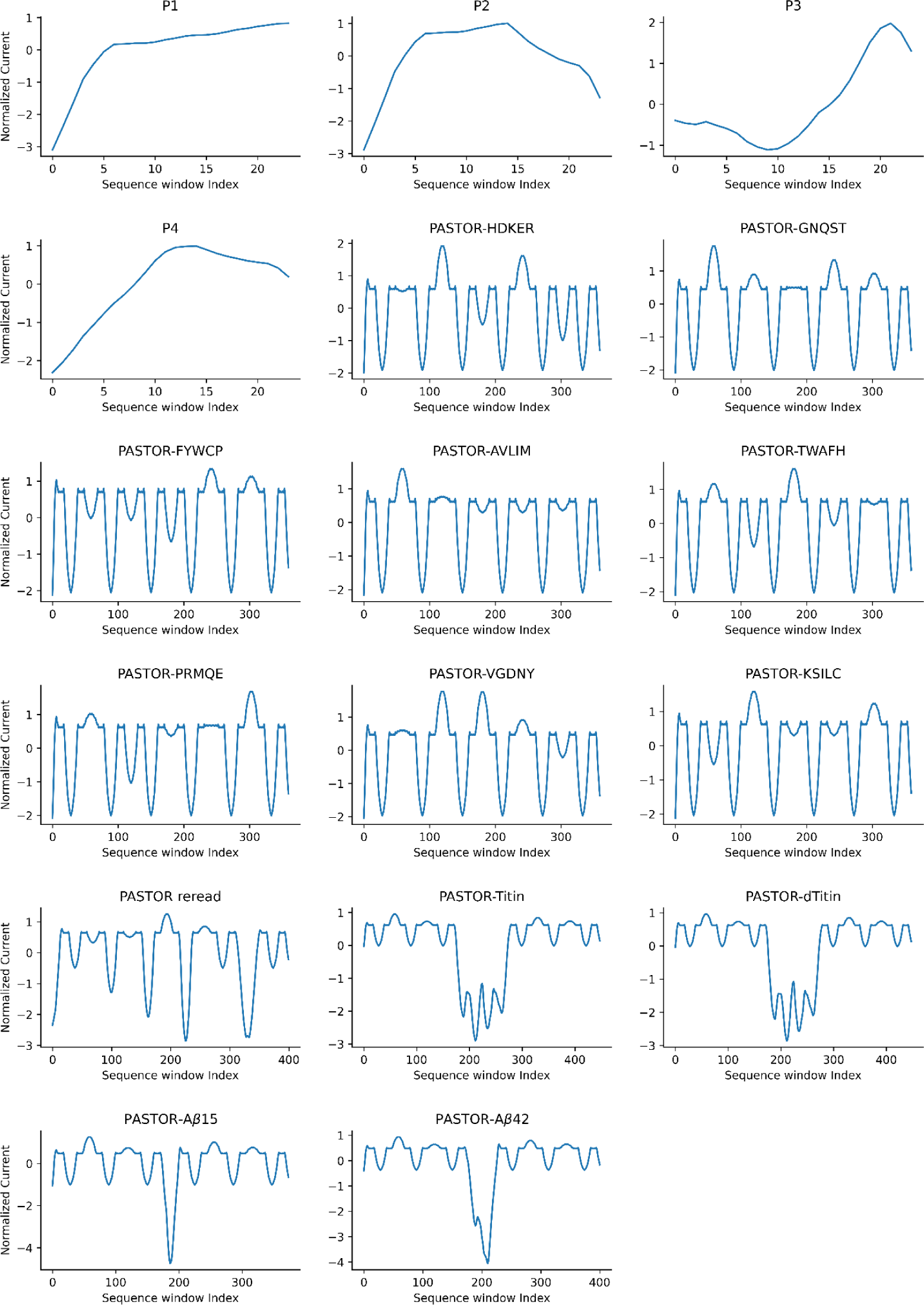
Normalized model traces generated by our biophysical model for all protein sequences analyzed in this study. Signal features contributed by protein domain unfolding are not included in the model.

**Supplementary Fig. 10.**
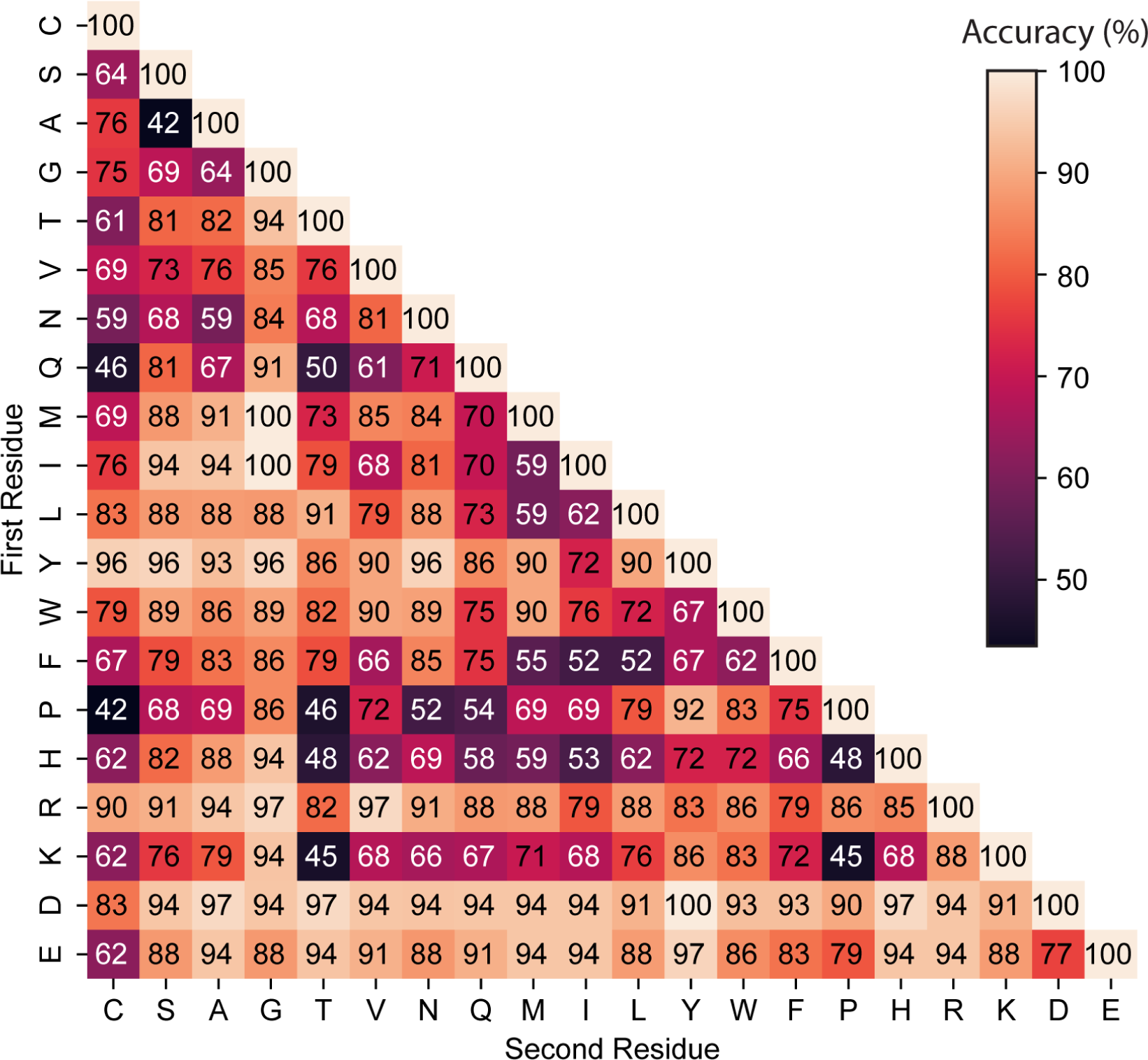
Heatmap each pairwise classification accuracy by a Random Forest model evaluated on the fixed test set, as in main Fig 4b, with values.

**Supplementary Fig. 11.**
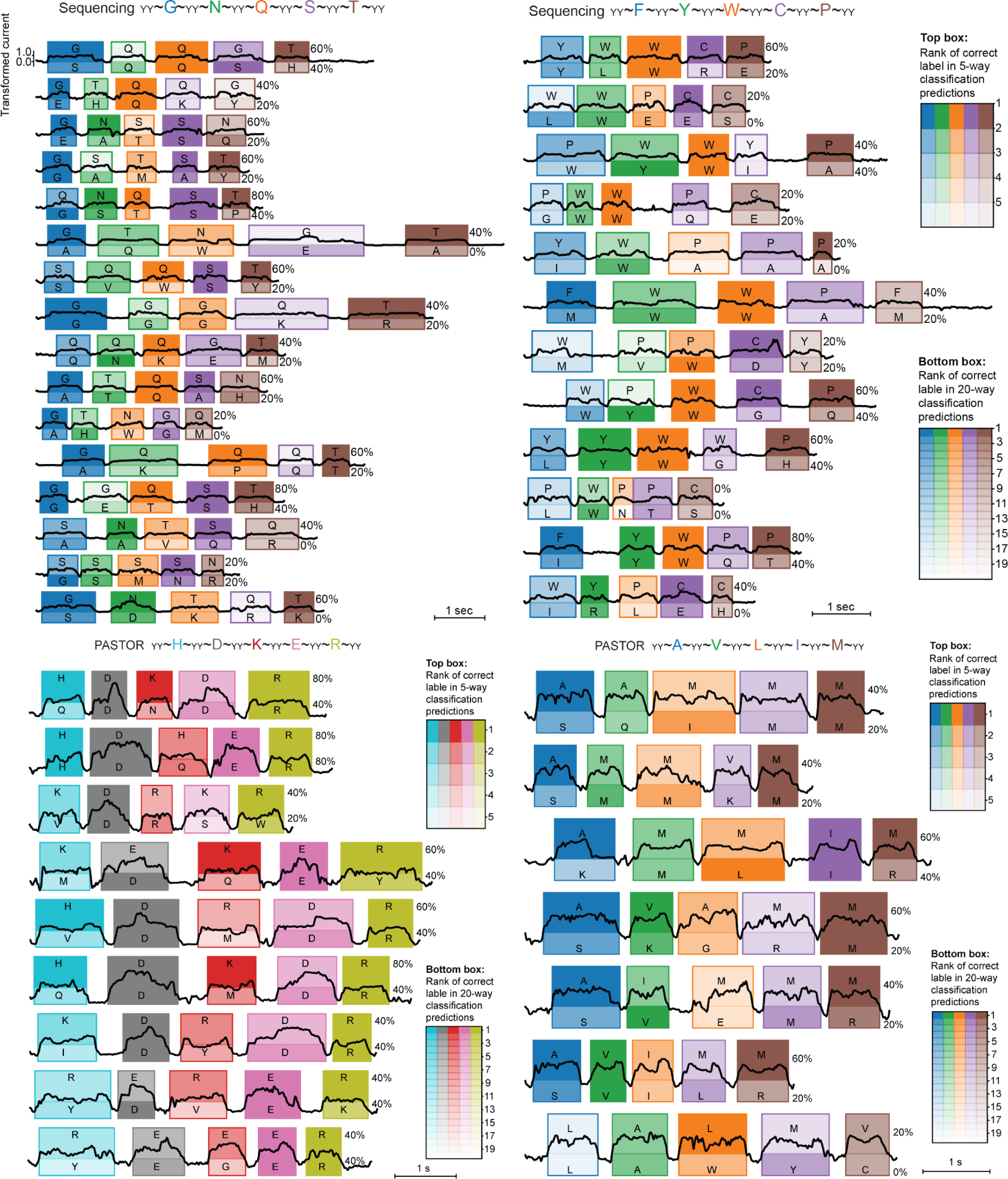
Reads of PASTOR-FYWCP, PASTOR-GNQST, and additional reads of PASTOR-HDKER and PASTOR-AVLIM. The traces of PASTOR-HDKER and PASTOR-AVLIM in Figure 2 were selected based on the 20-way classification model’s confidence of the predictions (confidence > 0.3 and > 0.17 for HDKER and AVLIM, respectively).

**Supplementary Fig. 12.**
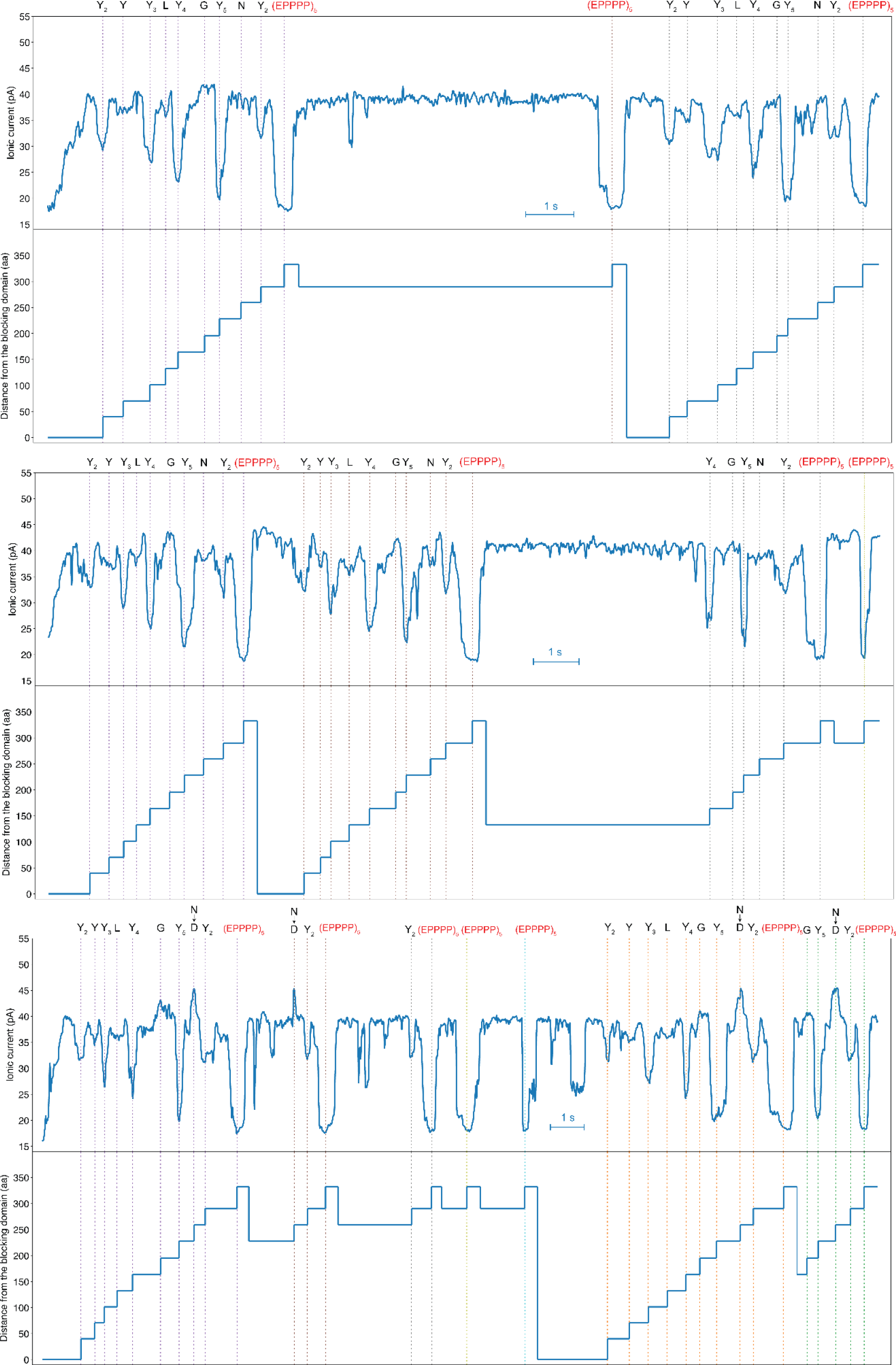
Example PASTOR-reread traces

**Supplementary Fig. 13.**
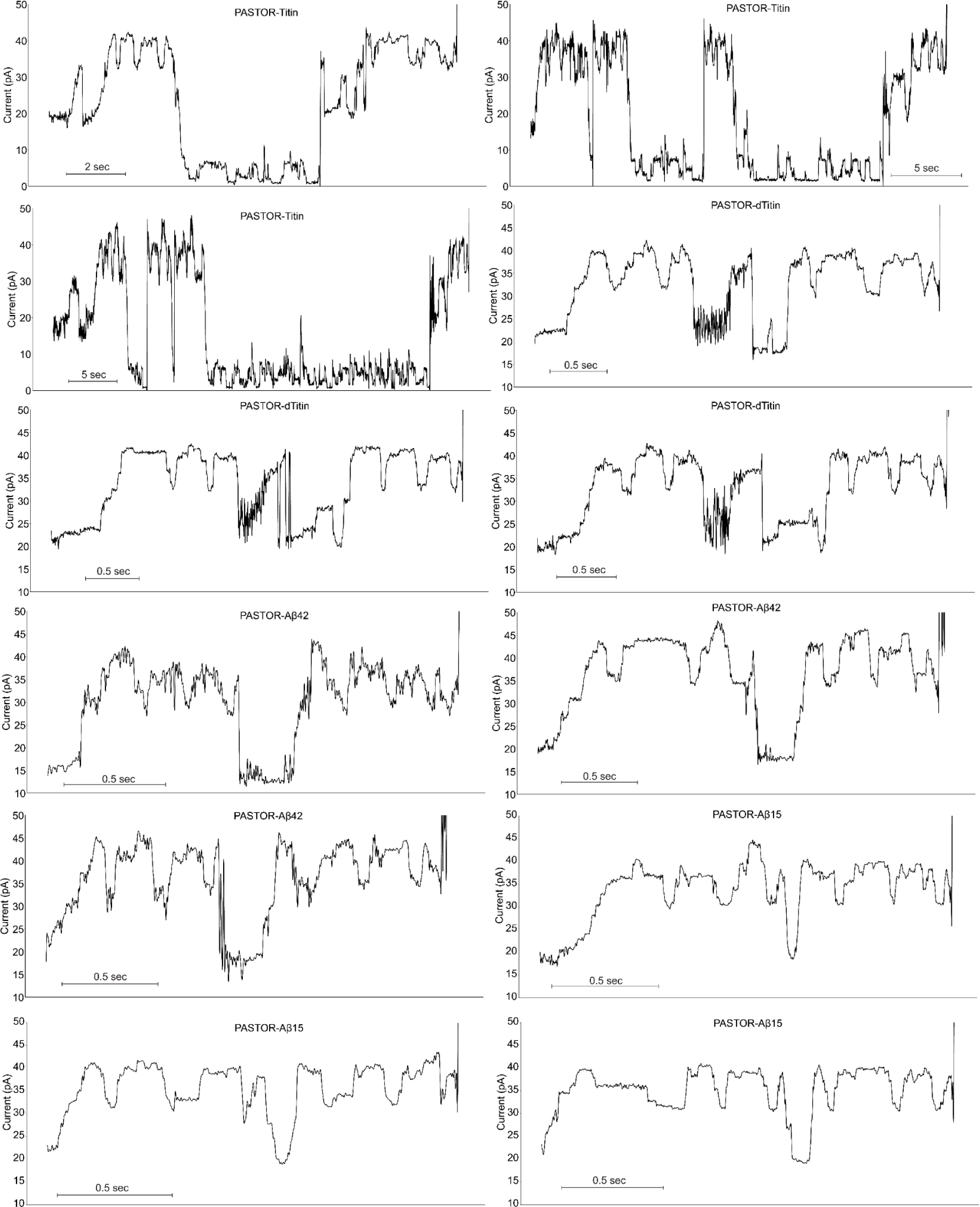
Filtered traces of example translocations of the PASTORs: -Titin, -dTitin, -Aβ42, and -Aβ15 proteins.

**Supplementary Fig. 14.**
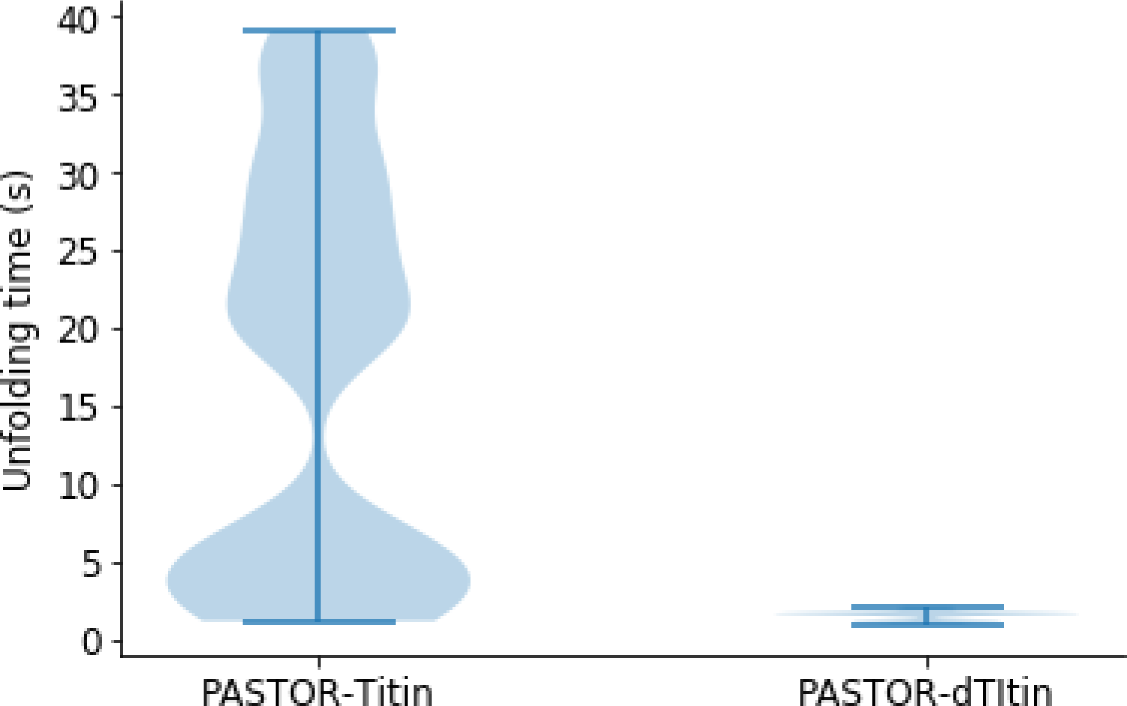
Distribution of total unfolding time for Titin (*n* = 21) and dTitin (*n* = 14).

**Supplementary Fig. 15.**
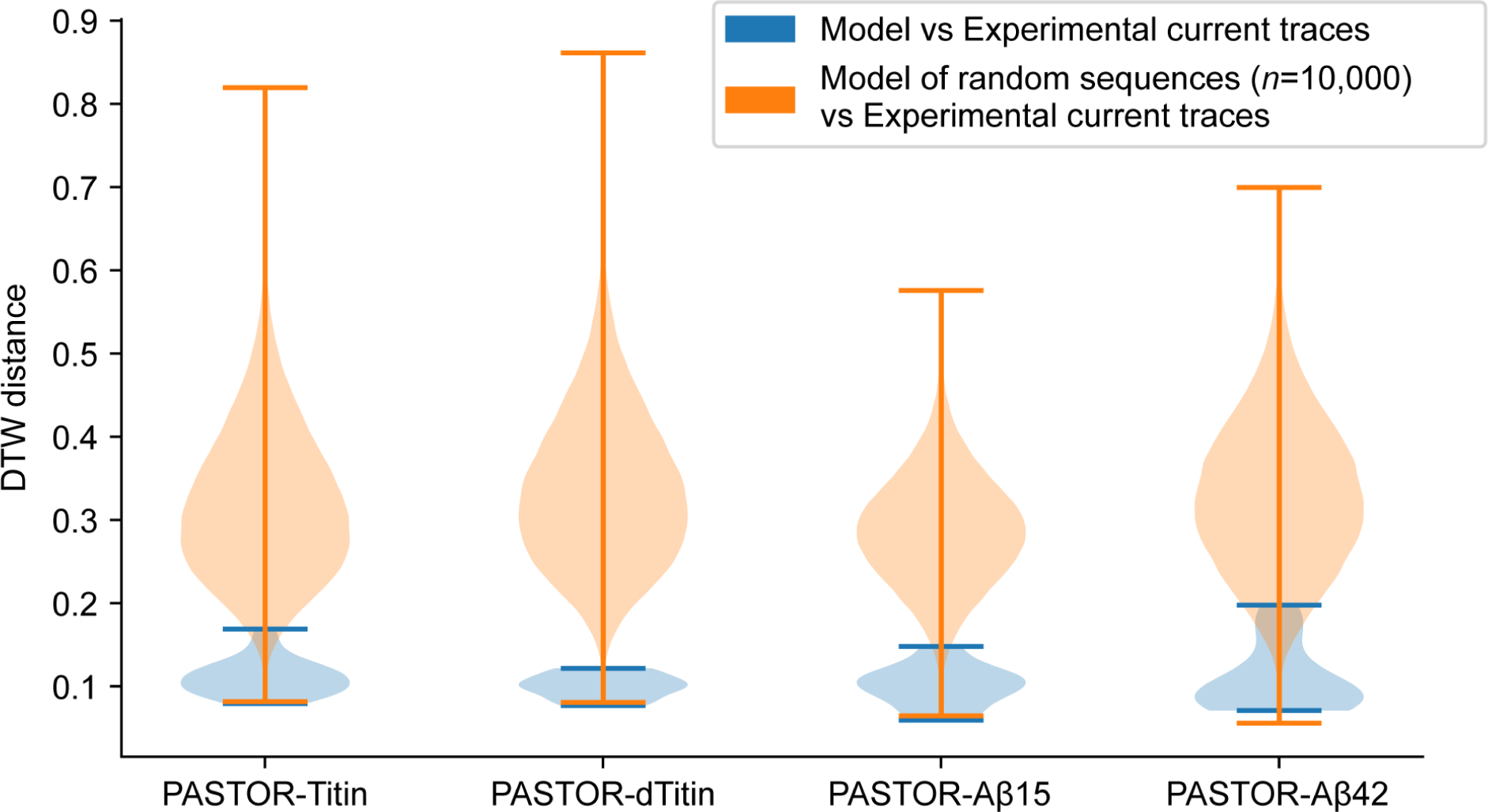
Distributions of the DTW distances of each of the protein translocations for folded domain proteins to model signal(s). In blue, they are compared to the model signal of the protein sequence, and in orange, they are each compared to the model of 10,000 random sequences derived from the same sequence distribution. The protein translocations include the regions corresponding to the folded domain translocation (state vii) and the N-terminal half of the PASTOR YY dips and VRs (state viii). The signals corresponding to the C-terminal half of the PASTOR YY dips and VRs (state v) and the folded domain unfolding (state vi) are excluded from the analysis, because the model does not predict unfolding patterns (main Figure 6b). *n* = 20, 12, 21, and 15 for PASTOR-Titin, PASTOR-dTitin, PASTOR-Aβ15, and PASTOR-Aβ42, respectively.

**Supplementary Table 1.** Set-wise classification accuracy. Accuracy of a hyperparameter-optimized Random Forest and CNN classifying VRs when the train and test set was composed only of VRs corresponding to the mentioned amino acids.

| Set Size | Random Forest Accuracy (%) | AAs |
| --- | --- | --- |
| 2 | 100 | Y, D |
| 3 | 95.0 | G, Y, D |
| 4 | 86.3 | A, W, R, D |
| 5 | 85.5 | G, V, W, R, D |
| 6 | 70.6 | C, G, L, Y, R, D |
| 7 | 65.9 | G, Q, W, F, R, D, E |
| 8 | 64.0 | A, T, I, Y, W, R, D, E |
| 9 | 59.0 | G, V, N, L, Y, W, R, D, E |
| 10 | 53.8 | A, G, V, N, Y, W, F, R, D, E |
| 11 | 50.0 | C, A, G, V, N, Y, W, F, R, D, E |
| 12 | 47.1 | C, A, G, T, V, Q, I, Y, W, R, D, E |
| 13 | 44.8 | A, G, T, V, N, Q, M, Y, W, F, R, D, E |
| 14 | 40.4 | C, A, G, T, V, N, Q, M, Y, W, F, R, D, E |
| 15 | 37.7 | C, S, G, T, V, N, Q, M, Y, W, F, R, K, D, E |
| 16 | 35.6 | C, A, G, T, V, N, Q, M, L, Y, W, F, R, K, D, E |
| 17 | 32.7 | C, S, A, G, T, V, N, Q, M, I, Y, W, F, H, R, D, E |
| 18 | 29.1 | C, S, A, G, T, N, M, I, L, Y, W, F, P, H, R, K, D, E |
| 19 | 28.9 | C, S, A, G, T, V, N, Q, M, I, L, Y, W, F, P, R, K, D, E |
| 20 | 28.3 | all 20 |

**Supplementary Table 2.**
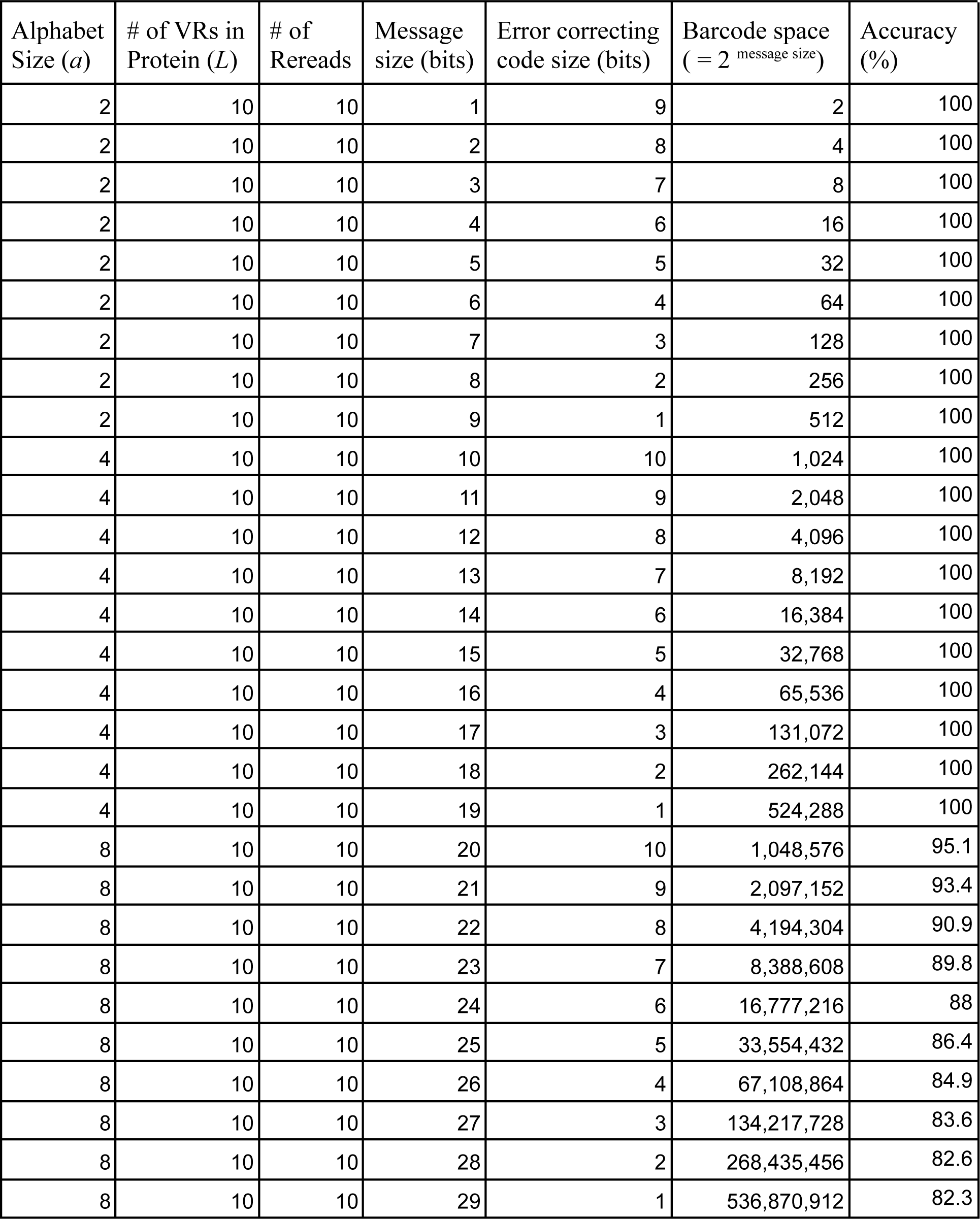
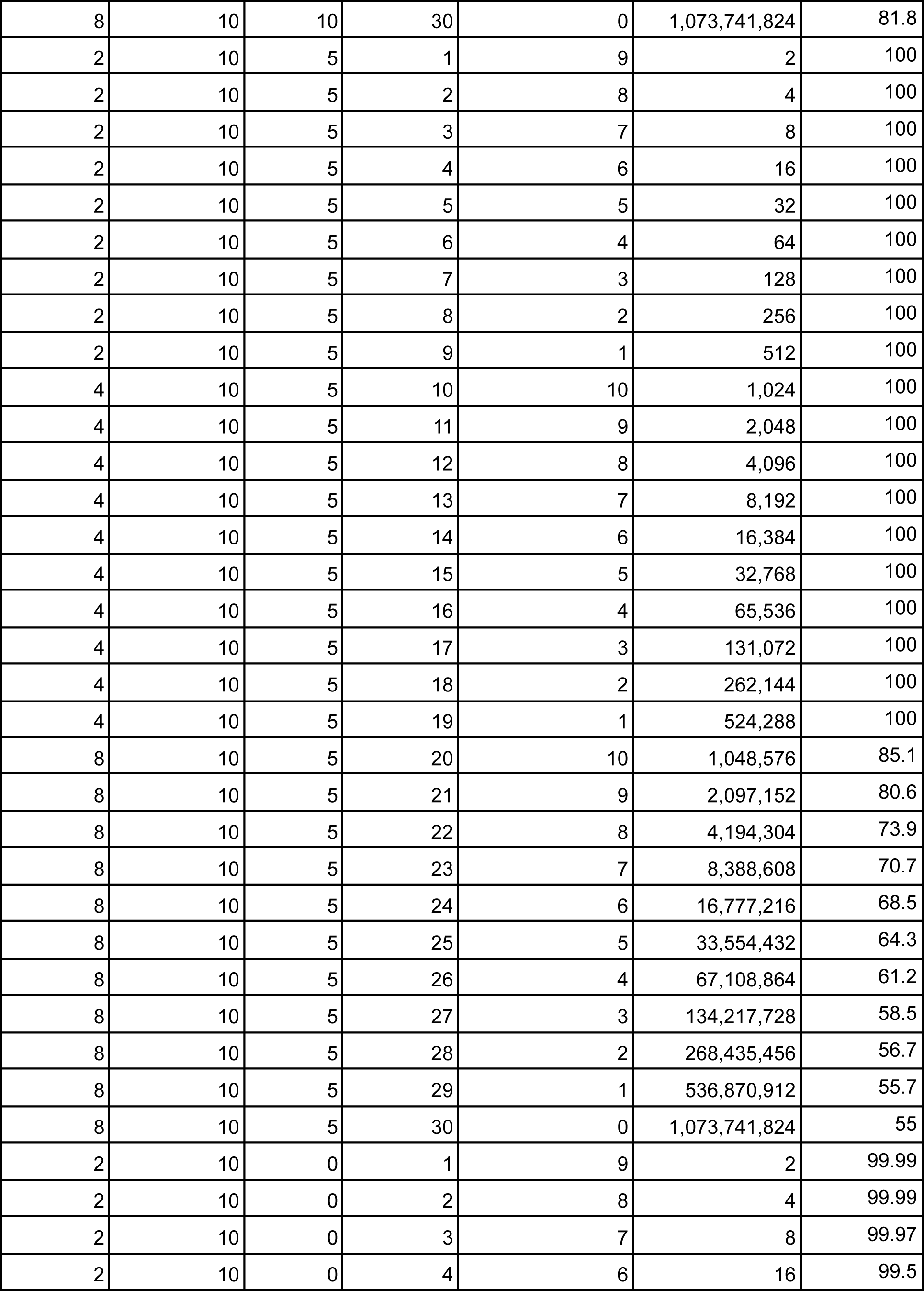
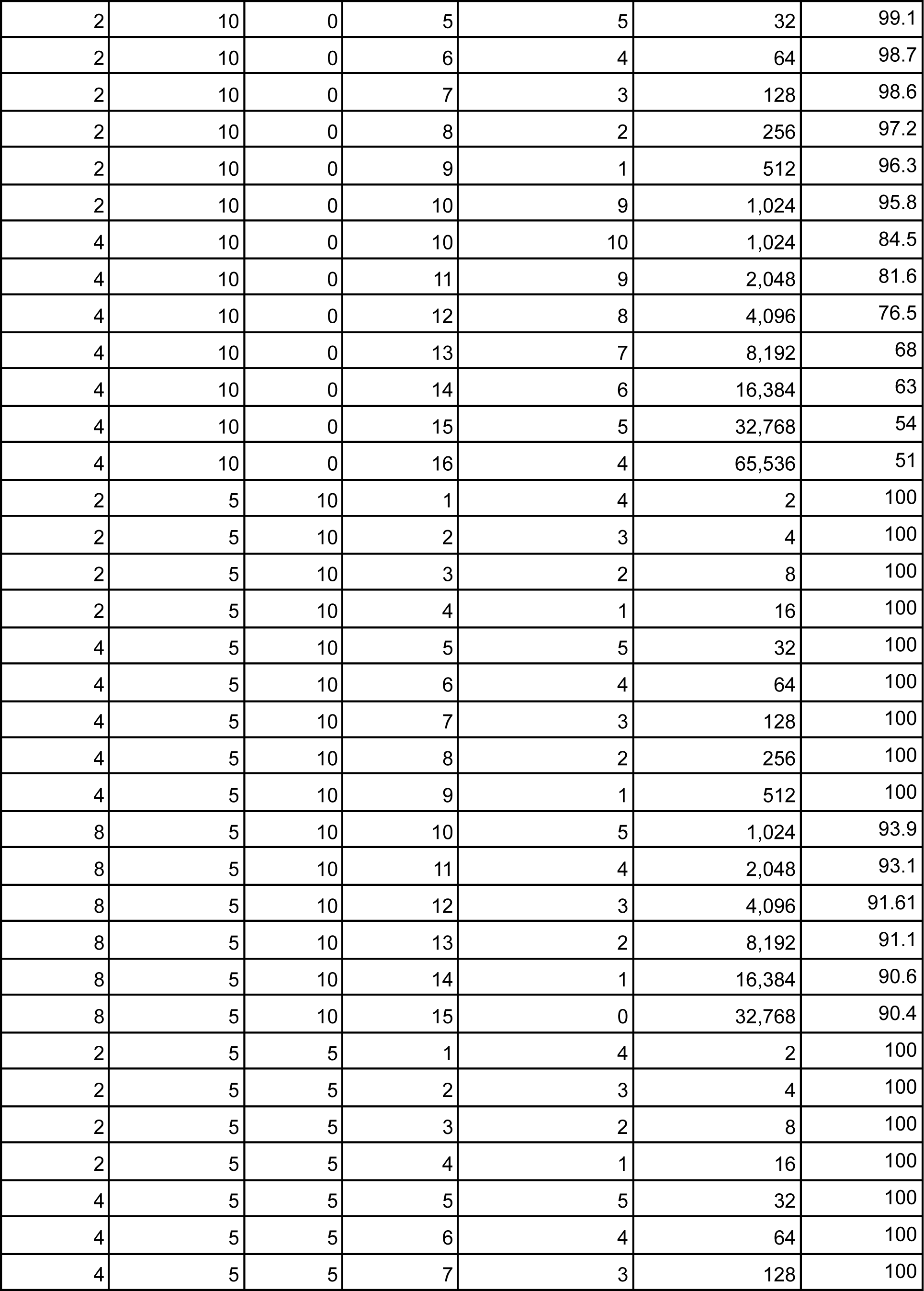
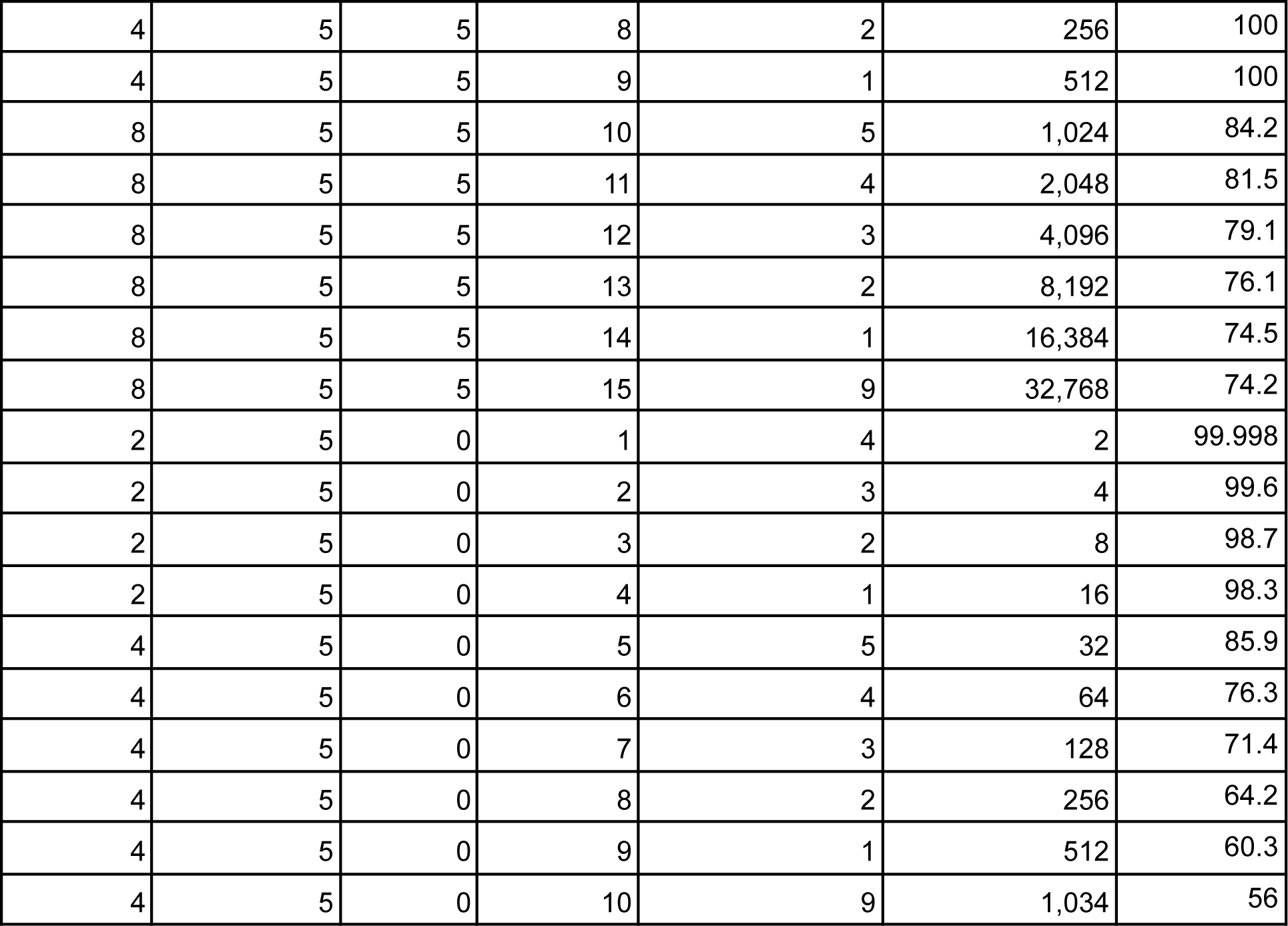
Bit allocation to error correcting codes in barcode design, as presented in Fig. 5g. Values are reported for the alphabet size that gives the highest possible accuracy for a given *L*, number of rereads, and message size. Conditions for which best accuracy is below 50% are excluded.

